# E proteins differentially co-operate with proneural bHLH transcription factors to sharpen neurogenesis

**DOI:** 10.1101/301093

**Authors:** Gwenvael Le Dréau, René Escalona, Raquel Fueyo, Antonio Herrera, Juan D. Martínez, Susana Usieto, Anghara Menendez, Sebastián Pons, Marian A. Martínez-Balbás, Elisa Martí

## Abstract

Basic HLH proteins heterodimerize with class I HLH/E proteins to promote transcription. Here we show that E proteins differentially co-operate with proneural bHLH transcription factors sharpening their neurogeneic activity. We find that inhibiting BMP signaling or its target ID2, in the chick embryo spinal cord, impairs the neuronal production from progenitors expressing ATOH1/ASCL1, but less severely that from progenitors expressing NEUROG1/2/PTF1a. We define the mechanisms of this differential response as a dual co-operation of E proteins with proneural proteins. E proteins synergize with bHLH proteins when acting on CAGSTG motifs, thereby facilitating the neurogenic activity of ASCL1/ATOH1 which preferentially bind to such motifs. Conversely, E proteins restrict the strong neurogenic potential of NEUROG1/2 by directly inhibiting their preferential binding to CADATG motifs. Since we find this mechanism to be conserved in corticogenesis, we propose this dual co-operation of E proteins with bHLH proteins as a novel though general feature of their mechanism of action.

## Introduction

The correct functioning of the vertebrate central nervous system (CNS) relies on the activity of a large variety of neurons that can be distinguished by their morphologies, physiological characteristics and anatomical locations (Zeng & Sanes, 2017). Such heterogeneity is generated during the phase of neurogenesis, once neural progenitors have been regionally specified and are instructed to exit the cell cycle and differentiate into discrete neuronal subtypes (Guillemot, 2007).

Neuronal differentiation and subtype specification are brought together by a small group of transcription factors (TFs) encoded by homologues of the *Drosophila* gene families *Atonal*, *Achaete*-*Scute*, *Neurogenins/dTap* and *p48/Ptf1a/Fer2* (Bertrand et al, 2002; Huang et al, 2014). These TFs represent a subgroup of the class II of helix-loop-helix proteins and all share a typical basic helix-loop-helix (bHLH) structural motif, where the basic domain mediates direct DNA binding to CANNTG sequences (known as E-boxes) and the HLH region is responsible for dimerization and protein-protein interactions (Bertrand et al, 2002; Massari & Murre, 2000). They are generally expressed in mutually exclusive populations of neural progenitors along the rostral-caudal and dorsal-ventral axes (Gowan et al, 2001; Lai et al, 2016). They are typically referred to as proneural proteins, since they are both necessary and sufficient to switch on the genetic programs that drive pan-neuronal differentiation and neuronal subtype specification during development (Guillemot, 2007). This unique characteristic is also illustrated by their ability to reprogram distinct neural and non-neural cell types into functional neurons (Masserdotti et al, 2016).

Regulating the activity of these proneural proteins is crucial to ensure the production of appropriate numbers of neurons without prematurely depleting the pools of neural progenitors. In cycling neural progenitors, the transcriptional repressors HES1 and HES5 act in response to Notch signalling to maintain proneural TF transcripts oscillating at low levels (Imayoshi & Kageyama, 2014). The proneural proteins are also regulated at the post-translational level. Ubiquitination and phosphorylation have been reported to control their stability, modify their DNA binding capacity or even terminate their transcriptional activity (Ali et al, 2011; Ali et al, 2014; Li et al, 2012; Quan et al, 2016). Furthermore, the activity of these proneural proteins is highly dependent on protein-protein interactions, and particularly on their dimerization status. It is generally admitted that these TFs must form heterodimers with the more broadly expressed class I HLH/E proteins in order to produce their transcriptional activity (Wang & Baker, 2015). In this way, the activity of proneural proteins can be controlled by upstream signals that regulate the relative availability of E proteins. Members of the Inhibitor of DNA binding (ID) family represent such regulators. As they lack the basic domain required for direct DNA-binding, ID proteins sequester E proteins through a physical interaction and thereby produce a dominant-negative effect on proneural proteins (Massari & Murre, 2000; Wang & Baker, 2015). Hence, several sophisticated regulatory mechanisms are available during development to control proneural protein activity and fine-tune neurogenesis.

Bone morphogenetic proteins (BMPs) contribute to multiple processes during the formation of the vertebrate CNS (Le Dreau & Marti, 2013; Liu & Niswander, 2005). Yet it is only in the past few years that their specific role in controlling vertebrate neurogenesis has begun to be defined (Choe et al, 2013; Le Dreau et al, 2012; Le Dreau et al, 2014; Segklia et al, 2012). During spinal cord development, SMAD1 and SMAD5, two canonical TFs of the BMP pathway (Massague et al, 2005), dictate the mode of division that spinal progenitors adopt during primary neurogenesis. Accordingly, strong SMAD1/5 activity promotes progenitor maintenance while weaker activity enables neurogenic divisions to occur (Le Dreau et al, 2014). This model explains how inhibition of BMP7 or SMAD1/5 activity provokes premature neuronal differentiation and the concomitant depletion of progenitors. However, it does not explain why the generation of distinct subtypes of dorsal interneurons are affected differently (Le Dreau et al, 2012), nor how BMP signaling affects the activity of the proneural proteins expressed in the corresponding progenitor domains.

Here, we have investigated these questions, extending our analysis to primary spinal neurogenesis along the whole dorsal-ventral axis. As such, we identified a striking correlation between the requirement of canonical BMP activity for the generation of a particular neuronal subtype and the proneural protein expressed in the corresponding progenitor domain. Inhibiting the activity of BMP7, SMAD1/5 or their downstream effector ID2 strongly impaired the production of neurons by spinal progenitors expressing either ATOH1 or ASCL1 alone, while it had a much weaker effect on the generation of the neuronal subtypes derived from progenitors expressing NEUROG1, NEUROG2 or PTF1a. We found that this differential responsiveness originates from a dual, E-box dependent mode of co-operation of the class I HLH/E proteins with the proneural proteins. E proteins interact with proneural proteins to aid their interaction with CAGSTG E-boxes, facilitating the ability of ASCL1 and ATOH1 to promote neurogenic divisions and hence, neuronal differentiation. Conversely, E proteins inhibit proneural protein binding to CADATG motifs, consequently restraining the ability of NEUROG1/2 that preferentially bind to these motifs to trigger neurogenic division and promote neuronal differentiation. Similar results were obtained in the context of corticogenesis, suggesting that this differential co-operation of E proteins with the distinct proneural proteins is a general feature of their mode of action.

## Results

### The canonical BMP pathway differentially regulates the generation of spinal neurons derived from progenitors expressing ASCL1/ATOH1 or NEUROG1/NEUROG2/PTF1a

We previously reported that BMP7 signalling through its canonical effectors SMAD1 and SMAD5, is differentially required for the generation of the distinct subtypes of dorsal spinal interneurons (Figure 1A and Le Dreau et al, 2012). Here, we extend this analysis to the generation of neuronal subtypes produced in the ventral part of the developing chick spinal cord. Inhibiting BMP7 or SMAD1/5 expression by *in ovo* electroporation of specific sh-RNA-encoding plasmids at stage HH14-15 produced a significant reduction in the generation of p2-derived Chx10^+^ (V2a) and Gata3^+^(V2b) subtypes 48 hours post-electroporation (hpe), whereas Evx1^+^ (V0v), En1^+^ (V1) interneurons and Isl1^+^ motoneurons were not significantly affected (Figure 1-figure supplement 1). These results revealed a correlation whereby the requirement of the canonical BMP pathway for the generation of discrete spinal neuron subtypes is linked to the proneural protein expressed in the corresponding progenitor domain (Figure 1B,C). The neuronal subtypes strongly affected by BMP7/SMAD1/5 inhibition (dI1, dI3, dI5: Figure 1B,C) were generated from spinal progenitors expressing ATOH1 (dP1) or ASCL1 alone (dP3, dP5). By contrast, all the neuronal subtypes deriving from spinal progenitors expressing either NEUROG1 alone (dP2, dP6-p1) or NEUROG2 (pMN) were much less severely affected (Figure 1B,C). Intriguingly, the V2a/b interneurons that display intermediate sensitivity to BMP7/SMAD1/5 inhibition are derived from p2 progenitors that express both ASCL1 and NEUROG1 (Misra et al, 2014), while the relatively insensitive dI4 interneurons are derived from dP4 progenitors that express PTF1a together with low levels of ASCL1 (Glasgow et al, 2005).

**Figure 1:**
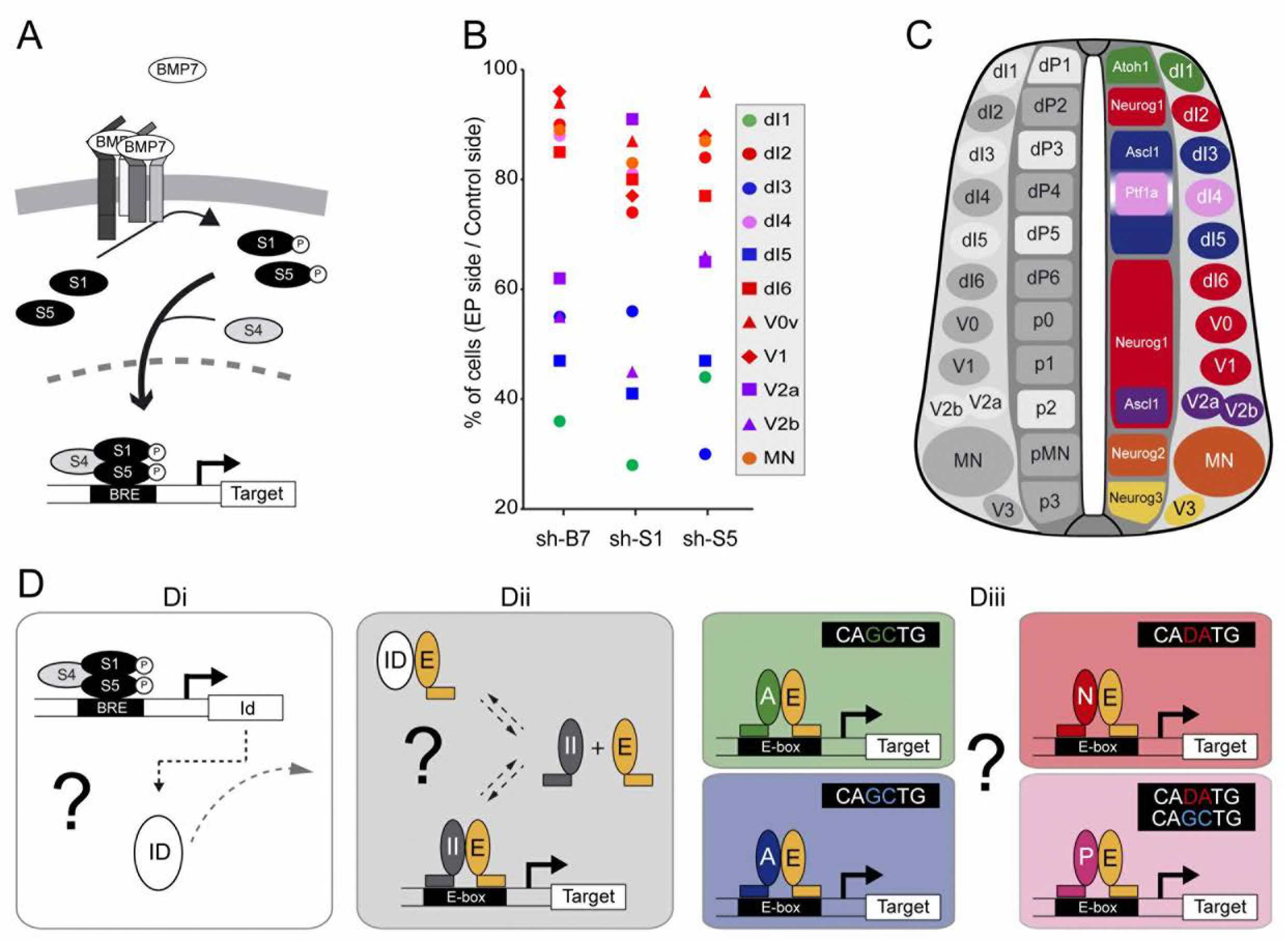
The canonical BMP pathway differentially regulates the generation of spinal neurons derived from progenitors that express ASCL1/ATOH1 or NEUROG1/NEUROG2/PTF1a. (A) Actors of the canonical BMP pathway (BMP7, SMAD1 and SMAD5) known to regulate spinal neurogenesis. (B) Dot-plot representing the spinal neuronal subtypes generated 48 hpe with plasmids producing sh-RNA targeting *cBmp7* (sh-B7), *cSmadl* (sh-S1) or *cSmad5* (sh-S5), comparing the electroporated side to the contra-lateral side. The colour code corresponds to the proneural proteins expressed in the corresponding progenitor domains, as shown in C. (C) Drawing of a transverse section of the developing spinal cord at mid-neurogenesis, highlighting: (left) the neuronal subtypes strongly (white) or moderately (grey) affected by inhibiting canonical BMP activity, and (right) a colour-coded representation of the proneural proteins expressed in the corresponding progenitor domains. (D) Working hypothesis whereby we propose to test if i) the canonical BMP activity is mediated by ID proteins; ii) ID proteins act by sequestering E proteins (E, orange), thereby inhibiting the activity of class II HLH/proneural proteins (II, grey); and iii) E proteins co-operate equally or differentially with the distinct proneural proteins as a function of their preferential binding to specific E-box sequences. The following figure supplement is available for figure 1: Figure supplement 1: Inhibiting the canonical BMP pathway affects the generation of ventral spinal neurons.

These correlations were particularly interesting in view of recent genome-wide ChIPseq studies that identified the optimal E-box (CANNTG) motifs bound by these TFs: ATOH1 and ASCL1 both preferentially bind to CAGCTG E-boxes (Borromeo et al, 2014; Castro et al, 2011; Klisch et al, 2011; Lai et al, 2011), whereas the optimal motif for NEUROGs is CADATG (where D stands for A, G or T: see Madelaine & Blader, 2011; Seo et al, 2007). Interestingly, most of the E-boxes bound by PTF1a in the developing spinal cord correspond to the CAGCTG motif favored by ASCL1 and ATOH1, yet PTF1a can bind to the NEUROG-like CAGATG motifs in a significant proportion of its targets genes (Borromeo et al, 2014). These observations suggested that the sensitivity of a given progenitor domain to canonical BMP activity originates from the intrinsic DNA-binding preferences of the different proneural bHLH TFs (Figure 1D). In many cell contexts, BMP signaling is mediated by ID proteins (Genander et al, 2014; Hollnagel et al, 1999; Moya et al, 2012), which physically sequester class I HLH/E proteins to produce a dominant-negative effect on proneural proteins (Figure 1D). While this hypothetical signaling cascade could explain the response of spinal progenitors expressing ASCL1 or ATOH1 to altered canonical BMP activity, it would not explain the relative insensitivity of the progenitors expressing NEUROG1, NEUROG2 or PTF1a. Therefore, we tested the veracity of these functional relationships to identify the basis of this differential response (Figure 1D).

### ID2 acts downstream of the canonical BMP pathway to differentially regulate the generation of spinal neurons derived from progenitors expressing ASCL1/ATOH1 or NEUROG1/NEUROG2/PTF1a

To test whether ID proteins are involved in canonical BMP signalling during spinal neurogenesis, we focused on ID2 (Figure 2A), not least because canonical BMP signalling is necessary and sufficient to promote *cId2* expression in the developing spinal cord (Figure 2-figure supplement 1 and Le Dreau et al, 2014). Moreover, *cId2* expression closely overlaps that described for the canonical BMP activity: restricted to the dorsal spinal cord early during patterning and later spreading ventrally within the ventricular zone during neurogenesis (Figure 2B-D and Le Dreau et al, 2012; Le Dreau et al, 2014). Inhibition of endogenous ID2 activity was triggered by *in ovo* electroporation of a sh-RNA specifically targeting chick *Id2* transcripts (sh-Id2, Figure 2-figure supplement 2A-E). This ID2 inhibition caused premature cell-autonomous differentiation at 48 hpe similar to that provoked by inhibiting SMAD1/5 (Figure 2E-K and Le Dreau et al, 2014). Conversely, overexpression of a murine ID2 construct reduced the proportion of electroporated cells that differentiated into neurons (Figure 2H,K and Figure 2-figure supplement 2F,G). ID2 overexpression could also partially impede the premature differentiation caused by both sh-Id2 and sh-Smad5 (Figure 2I-K). Similar results were obtained when measuring the activity of the pTubb3:luc reporter 24 hpe (Figure 2L).

**Figure 2:**
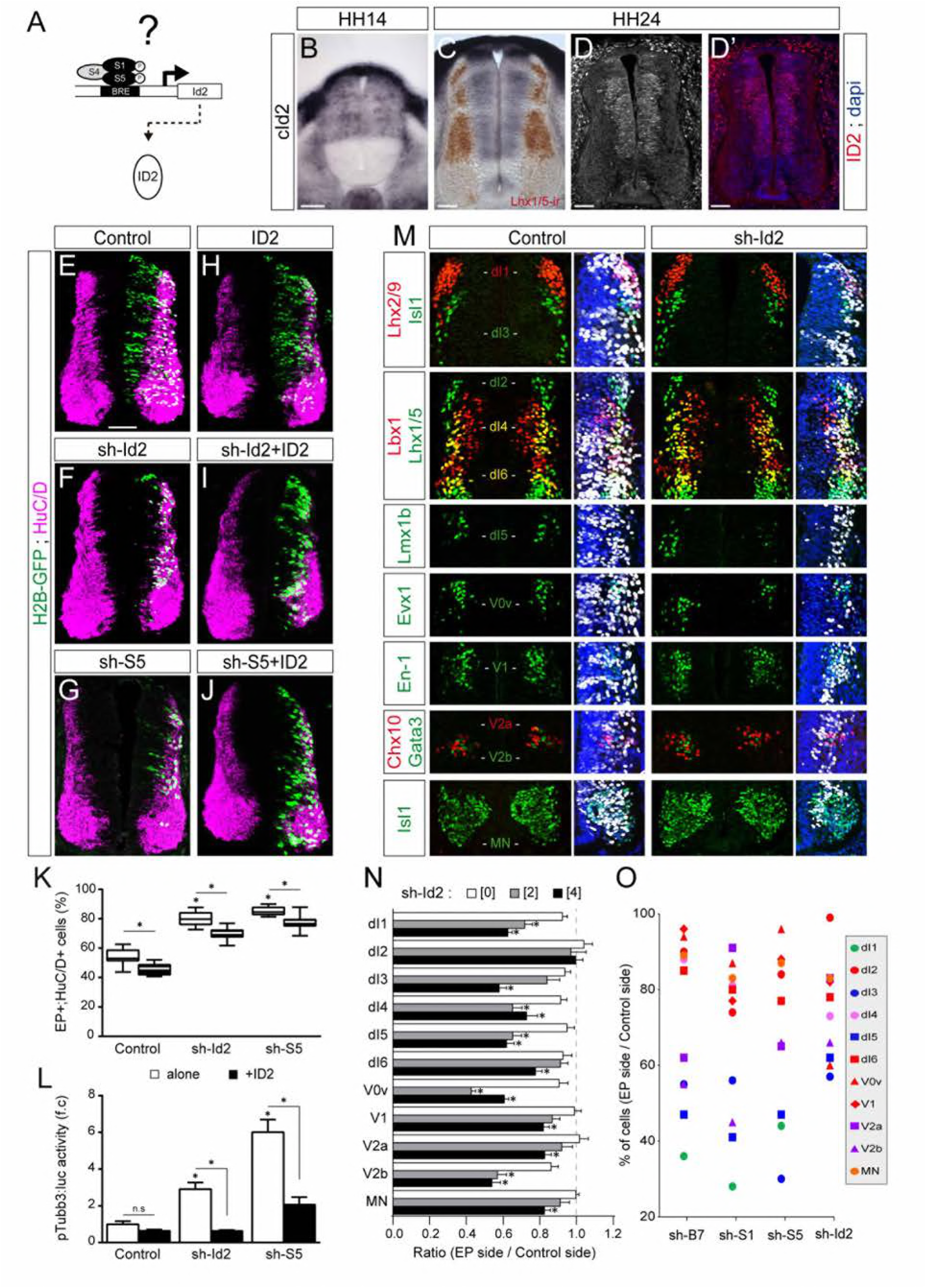
ID2 acts downstream of the canonical BMP pathway and it differentially regulates the generation of spinal neurons derived from progenitors expressing ASCL1/ATOH1 or NEUROG1/NEUROG2/PTF1a. (A) Hypothesis: I02 mediates the canonical BMP activity during spinal neurogenesis. (B, C) Detection of *cld2* transcripts by *in situ* hybridization in transverse spinal sections at stages HH14 (B) and HH24 (C), Lhx1/5 immunoreactivity (brown) was detected a *posteriori* (C). (D-D’) Endogenous clD2 immunoreactivity and DAPI staining at stage HH24. (E-J) Transverse spinal cord sections of electroporated cells (H2B-GFP+) that differentiated into neurons (HuCfD+) 48 hpe with: a control plasmid (E), plasmids producing sh-RNAs against c!d2 (sh-ld2, F) or cSmadS (sh-S5, G), a murine ID2 construct (H). and its combination with sh-ld2 (I) or sh-S5 (J). (K) Box-and-whisker plots obtained from n=7-l6 embryos: one-way ANOVA + Tukey’s test; ^*^P<0.05. (L) Activity of the pTubb3;luc reporter quantified 24 hpe in the conditions cited above, expressed as the mean fold change ± sem relative to the control, obtained from n=8-9 embryos; one-way ANOVA + Tukey’s test; ^*^P<0.05. (M) Representative images of the spinal neuron subtypes (identified with the combinations of the markers indicated) generated 48 hpe with control or sh-ld2. (N) Mean ratios ± sem or (O) dot-plots comparing the mean number of neurons on the electroporated and contralateral sides, obtained from n=8-11 embryos; one-way ANOVA + Tukey’s test; ^*^P<0.05. Scale bars, 50 μM. The following figure supplements are available for figure 2: Figure supplement 1: Regulation of ID2 expression by the canonical BMP pathway. Figure supplement 2; Modulation of ID2 activity in vivo.

We next analysed the consequences of ID2 inhibition on the generation of the different subtypes of spinal neurons, detecting a significant dose-dependent reduction in the generation of many neuronal subtypes (Figure 2M,N). The overall phenotype caused by ID2 inhibition was comparable to that triggered by inhibiting BMP7, SMAD1 or SMAD5: the neuronal subtypes deriving from progenitor domains expressing either ATOH1 or ASCL1 alone were globally more sensitive to ID2 inhibition than those derived from progenitor domains expressing NEUROG1, NEUROG2 or PTF1a (Figure 2O). Together, these results confirmed that ID2 acts downstream of the canonical BMP pathway in spinal neurogenesis and that it regulates distinctly the generation of spinal neurons derived from progenitors expressing ASCL1/ATOH1 and NEUROG1/NEUROG2.

### ID2 and E proteins counterbalance each other’s activity during spinal neurogenesis

We wondered whether ID2 contributes to spinal neurogenesis by sequestering E proteins (Figure 3A). Thus, we analysed the expression of these class I HLH genes during spinal neurogenesis. Transcripts from the *cTcf3/E2A* gene, which encodes the E12 or E47 alternative splice isoforms (Murre et al, 1989), were readily detected in the ventricular zone throughout the dorsal-ventral axis of the developing spinal cord, with apparently no domain-specific pattern (Figure 3B and Holmberg et al, 2008). Transcripts from the chicken HEB orthologue *cTcf12* were detected in the transition zone, following a dorsal-to-ventral gradient (Figure 3C). Previous studies reported that *E2*-*2* transcripts were barely detected in the developing murine spinal cord (Sobrado et al, 2009).

**Figure 3:**
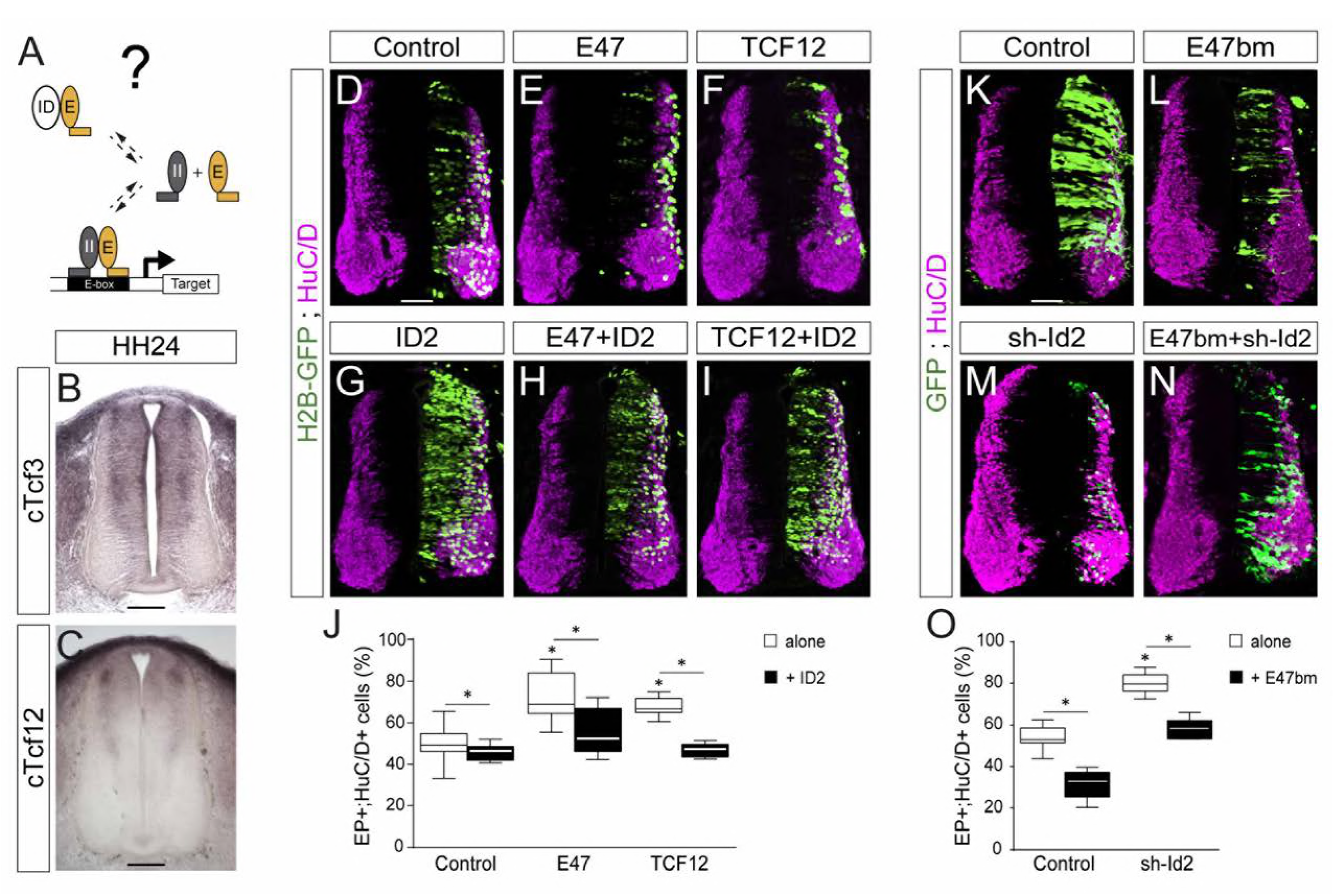
ID2 and E proteins counteract each other’s activity during spinal neurogenesis. (A) Hypothesis: ID2 sequesters E proteins during spinal neurogenesis. (B, C) Detection of *cTcf3/cE2a* (B) and *cTfc12* (C) transcripts by *in situ* hybridization in transverse spinal cord sections at stage HH24. (D-O) Transverse spinal cord sections of electroporated cells (GFP+ or H2B-GFP+) that differentiated into neurons (HuC/D+) 48 hpe with: a control (D), E47 (E), TCF12 (F), ID2 (G) or combinations of these (H, I); a control (K), E47bm (L), sh-ld2 (M) or their combination (N). (J, O) Box-and-whisker plots obtained from n=7-16 (J) and n=9-16 (O) embryos; one-way ANOVA + Tukey’s test; ^*^P<0 05 Scale bars, 50 μM The following figure supplement is available for figure 3: Figure supplement 1: E47bm rescues the premature neuronal differentiation caused by both E47 and TCF12.

The overexpression of E47 or TCF12 both produced a significant increase in the proportion of differentiated cells, a phenotype that was reverted by the concomitant electroporation of ID2 (Figure 3D-J). To inhibit the endogenous activity of E proteins, we took advantage of an E47 construct carrying mutations in its basic domain (E47bm) that acts in a dominant-negative manner over E proteins *in vivo* (Zhuang et al, 1998). Electroporation of E47bm inhibited neuronal differentiation in a cell autonomous manner, and it fully compensated for the premature differentiation caused by both E47 and TCF12 (Figure 3-figure supplement 1). The E47bm construct also rescued to a large extent the premature differentiation triggered by sh-Id2 (Figure 3K-O). Together, these results appear to confirm that the role played by ID2 during spinal neurogenesis depends on its ability to sequester E proteins.

### E47 co-operates distinctly with ASCL1/ATOH1 and NEUROG1/NEUROG2 to fine-tune neurogenic divisions during spinal neurogenesis

The results we obtained so far suggested that E proteins themselves might co-operate differently with the distinct proneural proteins during spinal neurogenesis (Figure 4A). To test this hypothesis, we first analyzed the consequences of expressing the mutant E47bm on the generation of spinal neuron subtypes. There was a marked reduction (≥ 50%) in the generation of Lhx2/9^+^ (dI1) and Tlx3^+^ (dI3/dI5) interneurons, which derive respectively from progenitors expressing ATOH1 and ASCL1 alone (Figure 4B,C,F). By contrast, electroporation of E47bm affected to a lesser extent (<25%) the generation of Lhx1/5+ (dI2/dI4/dI6-V1) or Isl1+ (MN) neuronal subtypes deriving from progenitors expressing NEUROG1 alone (dP2, dP6-V1), PTF1a (dP4) or NEUROG2 (pMN, Figure 4D-F). Hence, ATOH1 and ASCL1 appeared to be much more dependent on the activity of E proteins than NEUROG1, NEUROG2 and PTF1a to promote appropriate neuronal differentiation.

**Figure 4:**
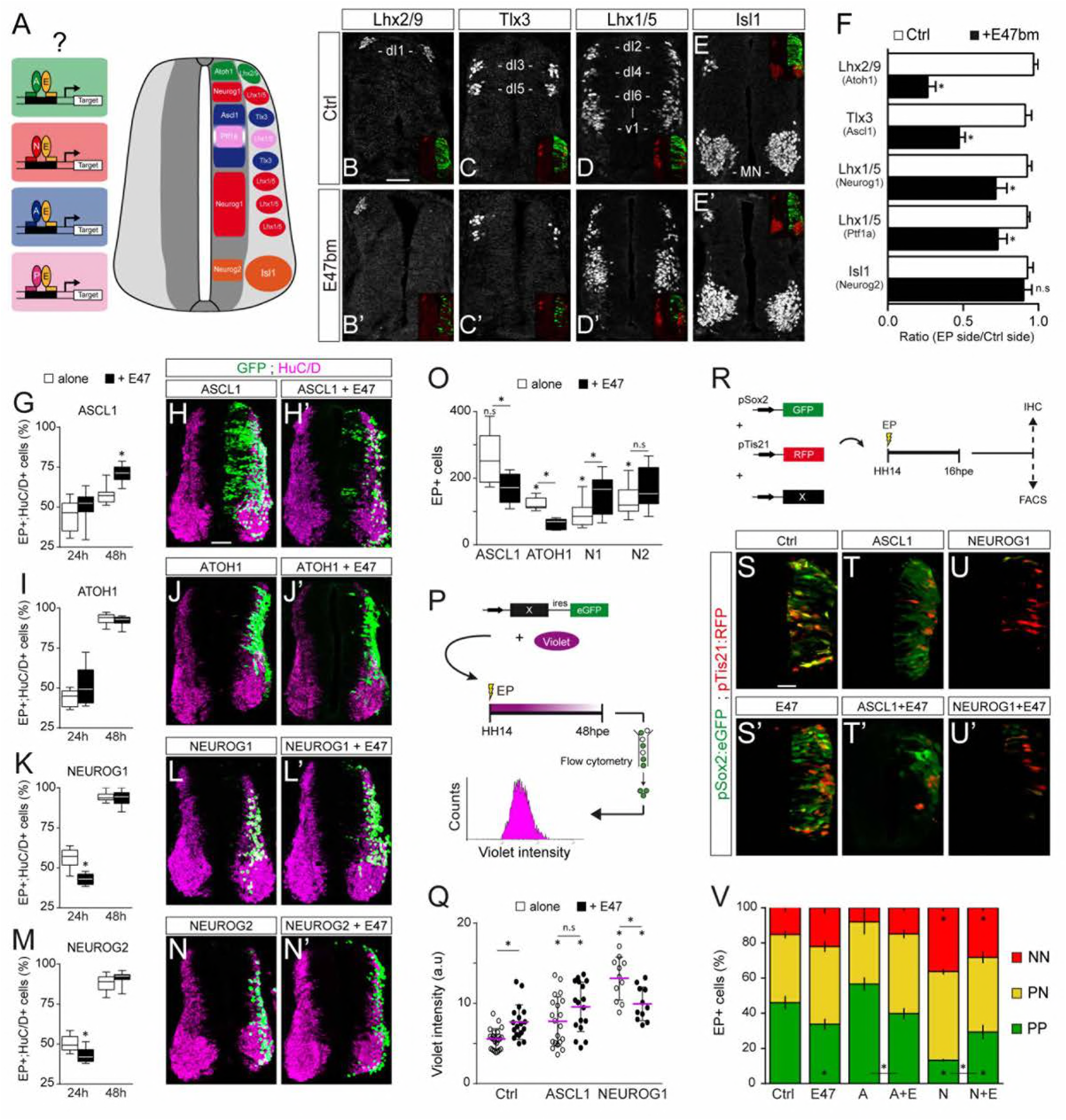
E47 co-operates with ASCL1/ATOH1 and NEUROG1/NEUROG2 distinctly to fine-tune neurogenic divisions during spinal neuro-genesis. (A) Hypothesis: E proteins co-operate differently with the distinct proneural proteins during spinal neurogenesis. (B-E) Representative images of spinal neurons expressing Lhx2/9 (dll, B-B’), Tlx3 (dl3/dl5, C-C’), Lhx1/5 (dl2/dl4/dl6-V1, D-D’) or Isl1 (MN, E-E’), 48 hpe with a control (B-E) or E47bm (B’-E’). (F)Mean ratios ±sem of neuron numbers on the electroporated side relative to the contralateral side, obtained from n=8-13 embryos; two-sided unpaired t-test: ^*^P<0.05. (G-N’) Transverse spinal cord sections of electroporated cells (GFP+ or H2B-GFP+) that differentiated into neurons (HuC/D^+^) 24 and 48 hpe with ASCL1 (G-H), ATOH1 (l-J). NEUROG1 (K-L) or NEUROG2 (M-N) alone (white) or together with E47 (black, H’-N). Box-and-whisker plots obtained from n=6-9 (G), 6-8 (I), 6-14 (K) and 7-12 (M) embryos; two-way ANOVA + Sidak’s test: ^*^P<0.05. (O) Mean number ± sem of electroporated cells quantified 48 hpe with the proneural proteins on their own (white) or together with E47 (black), obtained from 6-14 embryos: two-sided unpaired t-test; ^*^P<0 05. (P) Cell cycle exit assay. (Q) Mean Violet fluorescence intensity measured 48 hpe with a control, ASCL1 and NEUROG1 on their own (white) or together with E47 (black). The individual values (dots, n=11-23 embryos) and the mean (bars) are shown; one-way ANOVA + Tukey’s test and two-way ANOVA+Sidak s test; ^*^P<0.05. (R) Assessment of the modes of division of spinal progenitors. (S-U) Transverse spinal cord sections showing the activity of the pSox2:GFP and pTis21 :RFP reporters at 16 hpe, when electroporated in combination with control, ASCL1 or NEUROG1 on their own (S-U) or together with E47 (S’-U‘). (V) Mean proportion ± sem of cells identified as pSox2+/pTis21-(PP), pSox2 +/pTis21+ (PN) or pSox2-/pTis21 + (NN) when quantified by FACS, obtained from n=6-10 pools of embryos; two-way ANOVA + Tukey’s test; ^*^P<0.05. Scale bars, 50 μM. The following figure supplement is available for figure 4: Figure supplement 1: Effects of E47 and proneural proteins on spinal neuronal differentiation.

Next, we evaluated how E47 gain-of-function would modulate the neuronal differentiation induced when ASCL1, ATOH1, NEUROG1 and NEUROG2 are overexpressed (Figure 4G-O). From 24 hpe onwards, all 4 proneural bHLH proteins caused premature differentiation in a cell-autonomous and concentration dependent manner (Figure 4-figure supplement 1A-C). The presence of E47 accentuated the mild increase in neuronal differentiation provoked by ASCL1 at 24 hpe, and more significantly at 48 hpe (Figure 4G-H’). Accordingly, E47 provoked a significant reduction in the average number of electroporated cells generated 48 hpe of ASCL1 (Figure 4O). A similar tendency, albeit less pronounced, was observed when E47 was combined with ATOH1, especially in terms of the reduced average number of EP^+^ cells generated 48 hpe (Figure 4I-J’,O). Addition of E47 had the opposite effect when combined with NEUROG1 or NEUROG2: it significantly reduced the proportion of EP^+^;HuC/D^+^ cells obtained at 24 hpe and consequently increased the final numbers of EP^+^ cells observed at 48 hpe (Figure 4K-O). These results suggested that E47 differentially regulates the ability of ASCL1/ATOH1 and NEUROG1/NEUROG2 to promote cell cycle exit.

To assess cell cycle exit, a fluorescent cytoplasmic-retention dye that is only diluted on cell division was added at the time of electroporation and its mean fluorescence intensity was measured in FACS-sorted electroporated (GFP^+^) cells 48 hours later (Figure 4P). This assay demonstrated that E47 itself increased the mean Violet intensity, and further enhanced the mild increase caused by ASCL1 (Figure 4Q), indicating that E47 facilitates ASCL1’s ability to promote cell cycle exit. E47 had an opposite effect when combined with NEUROG1, significantly reducing the strong increase in Violet intensity caused by NEUROG1 (Figure 4Q), thereby confirming that E47 restricts NEUROG1’s ability to promote cell cycle exit.

We next studied how E47 influences the respective abilities of ASCL1 and NEUROG1 to regulate the balance between the 3 different modes of division that spinal progenitors can undergo during neurogenesis: symmetric proliferative divisions (PP), asymmetric divisions (PN), and symmetric neurogenic divisions (NN) (Le Dreau et al, 2014; Saade et al, 2013). To this end, we took advantage of the pSox2:eGFP and pTis21:RFP reporters that are specifically active during progenitor-generating (PP+PN) and neuron-generating (PN+NN) divisions, respectively (Saade et al, 2013). The effects of E47, ASCL1 and NEUROG1 on their activities were assayed 16 hpe by immunohistochemistry or quantified by FACS (Figure 4R). E47 caused a significant decrease in the proportion of pSox2:eGFP^+^;pTis21:RFP^-^ (PP) cells and a reciprocal increase in the proportion of pTis21:RFP^+^ (PN+NN) neurogenic divisions relative to the controls (Figure 4S,S’,V). While we did not detect any significant change in the proportions of PP, PN and NN cells in response to ASCL1 alone in these conditions, we did observe an increase in neurogenic divisions at the expense of proliferative divisions when ASCL1 was combined with E47 (Figure 4T,T’,V). Conversely, E47 significantly restrained NEUROG1’s ability to trigger neurogenic divisions at the expense of PP divisions (Figure 4U,V). Assessing the activity of the pSox2:luc reporter confirmed these results, further showing that E47 facilitates the ability of both ASCL1 and ATOH1 to repress pSox2 activity, whereas it restricts the repressive effects of both NEUROG1 and NEUROG2 (Figure 4-figure supplement 1D). Together, these results revealed that E47 co-operates distinctly with ASCL1/ATOH1 and NEUROG1/NEUROG2 to fine-tune neurogenic divisions during spinal neurogenesis.

### E47 co-operates distinctly with ASCL1 and NEUROG1 in an E-box dependent manner and through physical interactions

To identify the molecular mechanisms underlying the differential co-operation of E proteins with the distinct proneural proteins, we focused on the interaction of E47 with ASCL1 and NEUROG1. A DNA-binding deficient version of NEUROG1 (NEUROG1-AQ, (Sun et al, 2001), was unable to transactivate the NEUROG-responsive pNeuroD:luc reporter or to promote neuronal differentiation (Figure 5-figure supplement 1). Hence, the ability of NEUROG1 to trigger neuronal differentiation during spinal neurogenesis depends on its transcriptional activity, as previously reported for ASCL1 and ATOH1 (Nakada et al, 2004).

Genome-wide ChIP-seq studies have established that the preferential E-box motifs bound by ASCL1, E47 and NEUROG1 correspond respectively to CAGCTG (Borromeo et al, 2014; Castro et al, 2011), CAGSTG (where S stands for C or G: Lin et al, 2010; Pfurr et al, 2017) and CADATG (where D stands for A, G or T: Madelaine & Blader, 2011; Seo et al, 2007). In the light of these intrinsic preferences, we tested how E47 modulates the abilities of ASCL1 and NEUROG1 to bind to DNA and activate transcription via different E-boxes (Figure 5A and Figure 5-figure supplement 2A). E47 acted in synergy with both ASCL1 and NEUROG1 to drive transcription of the pkE7:luc reporter under the control of 7 CAGGTG repeats (Figure 5B,C). By contrast, E47 and ASCL1 only weakly transactivated the pNeuroD:luc reporter, the promoter of which contains 9 CADATG E-boxes and 1 CAGGTG box (Figure 5D). A similar result was obtained with a version of the pNeuroD:luc reporter in which the single CAGGTG motif was destroyed by mutagenesis (Figure 5-figure supplement 2B), reinforcing the idea that both E47 and ASCL1 preferentially bind to CAGSTG sequences. Intriguingly, E47 markedly reduced the ability of NEUROG1 to enhance pNeuroD:luc activity (Figure 5E). A similar result was obtained with the mutated version of the pNeuroD:luc reporter (Figure 5-figure supplement 2C), ruling out the possibility that specific E47 binding to the single CAGGTG motif in this promoter caused this inhibition. *In vitro* ChIP assays demonstrated that E47 can bind to and enhance ASCL1 binding at the 7 CAGGTG-containing promoter region of the pkE7:luc reporter (Figure 5F), consistent with the notion that their heterodimerization is required for optimal binding and subsequent transcriptional activation. By contrast, E47 caused a significant reduction in the amount of NEUROG1 bound to the promoter region of the pNeuroD:luc reporter (Figure 5G). The fact that E47 itself bound to this promoter region suggested that E47 and NEUROG1 compete for binding to CADATG motifs (Figure 5G), although E47 cannot transactivate them as potently as NEUROG1 (Figure 5E). Together, these results revealed that E47 acts in synergy with both ASCL1 and NEUROG1 when binding to its own optimal E-box (CAGSTG), while it somehow impedes NEUROG1 from binding to CADATG motifs.

**Figure 5:**
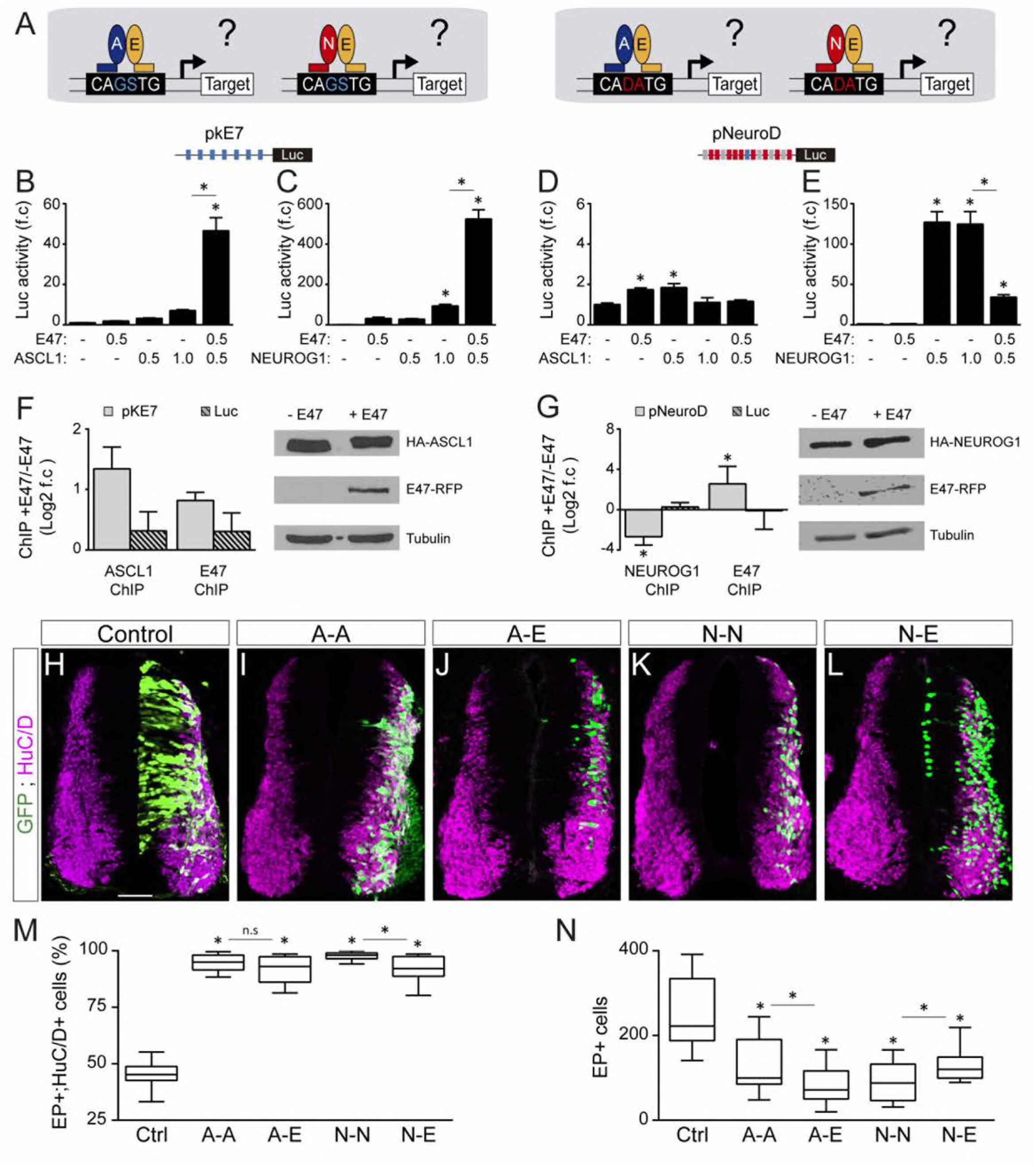
E47 co-operates with ASCL1 and NEUROG1 in an E-box dependent manner and through physical interactions. (A) Hypothesis: E proteins co-operate differently with proneura! proteins in different E-box contexts. (B-E) Activity of the pkE7 (B, C) and pNeuroD (D. E) luciferase reporters measured 24 hpe with a control, E47 and ASCL1 (B, D) or NEUROG1 (C, E), expressed as the mean fold change ± sem relative to the control, obtained from n=8 embryos; one-way ANOVA + Tukey’stest; ^*^P<0.05. (F-G) ChIP assays performed on the pkE7 (F) or pNeu-roD (G) promoter regions (light grey), or luciferase ORF (Luc, striped grey), in HEK293 cells 24 hours after transfection with HA-ASCL1 (F) or HA-NEUROG1 (G) on their own or together with E47-RFP expressed as Log2 values of the mean fold change ± sem in DNA binding measured in the presence of E47 relative to absence of E47, obtained from n=3 (F) or n=5 (G) experiments; two-sided one sample t-test: ^*^P<0 05 The HA-ASCL1, HA-NEUROG1 and E47-RFP proteins probed in Western blots, with Tubulin-beta as a transfection control. (H-L) Transverse spinal cord sections of electroporated cells (GFP+) that differentiated into neurons (HuC/D+) 48 hpe with a control (H), ASCL1 or NEUROG1 homodimer (A-A, I; N-N, K), or ASCL1-E47 or NEUROG1-E47 heterodimers (A-E, J; N-E, L). (M) Box-and-whisker plots obtained from n=12-15 embryos; one-way ANOVA + Tukey’s test: ^*^P<0.05. (N) Mean number of electroporated cells (GFP+) generated 48 hpe in the conditions cited above, calculated from n=11-14 embryos; one-way ANOVA + Tukey’s test; ^*^P<0.05. Scale bars. 50 μM. The following figure supplements are available for figure 5: Figure supplement 1: The ability of NEUROG1 to induce spinal neuronal differentiation depends on its DNA-binding. Figure supplement 2: E-box dependent activity of E47, ASCL1 and NEUROG1 during spinal neurogenesis. Figure supplement 3: Characterization of the tethered constructs of bHLH dimers.

To confirm that the differential co-operation of E47 with ASCL1 and NEUROG1 is due to a direct physical interaction, we compared the activity of tethered constructs that were designed to produce homodimers of ASCL1 (A-A) and NEUROG1 (N-N), or heterodimers with E47 (A-E, N-E: Figure 5-figure supplement 3A-C). Consistent with the results obtained with monomers, A-E heterodimers were significantly more potent than A-A homodimers in driving pkE7:luc activity (Figure 5-figure supplement 3D), while N-N homodimers transactivated pNeuroD:luc much more strongly than N-E heterodimers (Figure 5-figure supplement 3E). A-A and A-E promoted similar neuronal differentiation 48 hpe (Figure 5H-J,M), but the average number of EP^+^ cells obtained after A-E electroporation was significantly less than after A-A electroporation (Figure 5N), suggesting that A-E promotes early neurogenic divisions more potently than A-A. This idea was supported by the ability of A-E to repress pSox2:luc activity at 20 hpe, unlike A-A (Figure 5-figure supplement 3F). As for NEUROG1, N-N was significantly more potent than N-E at promoting neuronal differentiation (Figure 5K-M), at reducing the average number of EP^+^ cells generated 48 hpe (Figure 5N) and at repressing pSox2:luc activity (Figure 5-figure supplement 3F). Thus, the tethered constructs performed like the monomers (Figure 4G-V), supporting the conclusion that E47 facilitates the ability of ASCL1 and restrains that of NEUROG1 to trigger neurogenic divisions during spinal neurogenesis.

### E47 co-operates differentially with ASCL1 and NEUROG1/NEUROG2 during corticogenesis

We were interested to determine if this differential co-operation of E47 with the distinct proneural proteins could be extended to other regions of the developing CNS. We tested this hypothesis in the developing cerebral cortex, as NEUROG1/2 and ASCL1 all contribute to neurogenesis in this region in mammals (Huang et al, 2014). The development of the cerebral cortex in birds actually shows unexpected similarities to mammalian corticogenesis, including the conservation of its temporal sequence of neurogenesis (Dugas-Ford et al, 2012; Suzuki et al, 2012). As in mammals, corticogenesis in the chick embryo originates from a region of the dorsal pallium expressing PAX6 (Figure 6A,B and Suzuki et al, 2012). From E3 to E5, an early phase of corticogenesis produces the first SOX2^-^;HuC/D^+^ cortical neurons, which are generated specifically from PAX6^+^;TBR2^-^ radial glia-like progenitors that divide at the apical surface, as in mammals (Figure 6C-D). Cortical TBR2^+^ progenitors that divide basally, similar to mammalian intermediate progenitor cells, appear at around E5 (Figure 6D-D”). The cortical neurons produced during this early phase express TBR1 (Figure 6-figure supplement 1A), as well as other markers typically expressed by mammalian deep-layer neurons (Dugas-Ford et al, 2012; Suzuki et al, 2012). At E4, *cNeurog1* and *cNeurog2* transcripts were detected in a salt-and-pepper fashion in the cortical PAX6^+^ region (Figure 6E,F), whereas *cAscl1* expression was detected strongly in the sub-pallium and more weakly in the developing cerebral cortex (Figure 6G). These expression patterns seen in early chicken embryos are very similar to what is observed in the developing mammalian telencephalon (Huang et al, 2014).

**Figure 6:**
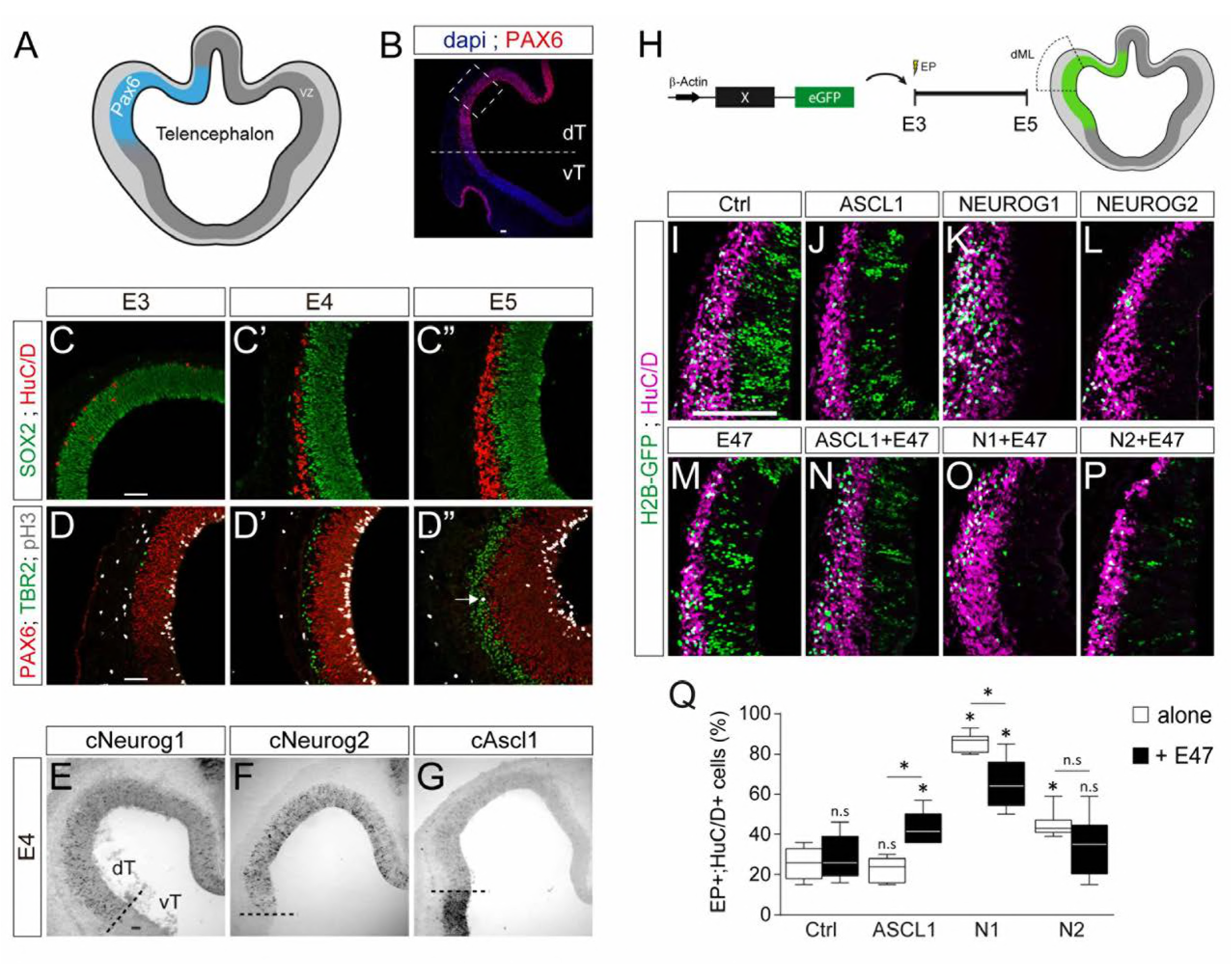
E47 co-operates differentially with ASCL1 and NEUROG1/NEUROG2 during corticogenesis. (A) Scheme of the embryonic chick telencephalon at early stages of neurogenesis. (B-D) Coronal telencephalic sections showing PAX6 immuno-reactivity and the cell nuclei (DAPI) at low magnification at E4 (B), cortical progenitors and differentiating neurons (SOX2+ and HuC/D+, C), apical progenitors (PAX6+;TBR2-, D) and mitotic basal progenitors (TBR2+;pH3+, arrow in D”) at E3 (C, D), E4 (C’, D’) and E5 (C”, D”). (E-G) Detection of *cNeurogl* (E), *cNeurog2* (F) and *cAscl1* (G) transcripts by *in situ* hybridization at E4. (H) *In ovo* electroporation of the chick telencephalon. (I-P) Coronal telencephalic sections of electroporated cells (GFP+) that differentiated into neurons (HuC/D+) 48 hpe with a control (I), ASCL1 (J), N EUROG1 (K) or NEUROG2 (L) on their own or together with E47 (M-P). (Q) Box-and-whisker plots obtained from n=5-9 embryos; one-way ANOVA + Tukey’s test; ^*^P<0.05. Scale bars, 50 μM. dT/vT, dorsal and ventral telencephalon; dML, dorso-medial-lateral; VZ, ventricular zone. The following figure supplement is available for figure 6: Figure supplement 1: Neurogenesis and concentration dependent effects of proneural proteins during early chick corticogenesis.

To test how E47 modulates the activity of ASCL1 and NEUROG1/2 in the developing chick cerebral cortex, we electroporated the dorsal telencephalic region at E3 *in ovo* and analysed neuronal differentiation 2 days later (Figure 6H). Both NEUROG1 and NEUROG2 triggered significant neuronal differentiation in the developing cerebral cortex in a cell autonomous and dose-dependent manner, whereas ASCL1 overexpression had only a minor effect *per se* (Figure 6I-L and Figure 6-figure supplement 1B). E47, which itself had no obvious effect at this concentration (Figure 6I,M,Q), significantly increased neuronal differentiation when combined with ASCL1 (Figure 6J,N,Q). Conversely, E47 markedly reduced the ability of NEUROG1, and to a lesser extent that of NEUROG2, to promote neuronal differentiation (Figure 6K-Q). These results suggest that E47 also co-operates differentially with ASCL1 and NEUROG1/2 in the context of cortical neurogenesis.

## Discussion

Class I HLH/E proteins are generally described as obligatory and permissive co-factors for proneural proteins, which must form heterodimers to become active and regulate transcription (Wang & Baker, 2015). The main findings of this study are that the co-operation between E proteins and proneural proteins might be more complex than originally thought. Indeed, our results revealed that E proteins can facilitate or restrain the transcriptional activity of proneural bHLH TFs depending on the E-boxes involved (Figure 7).

**Figure 7:**
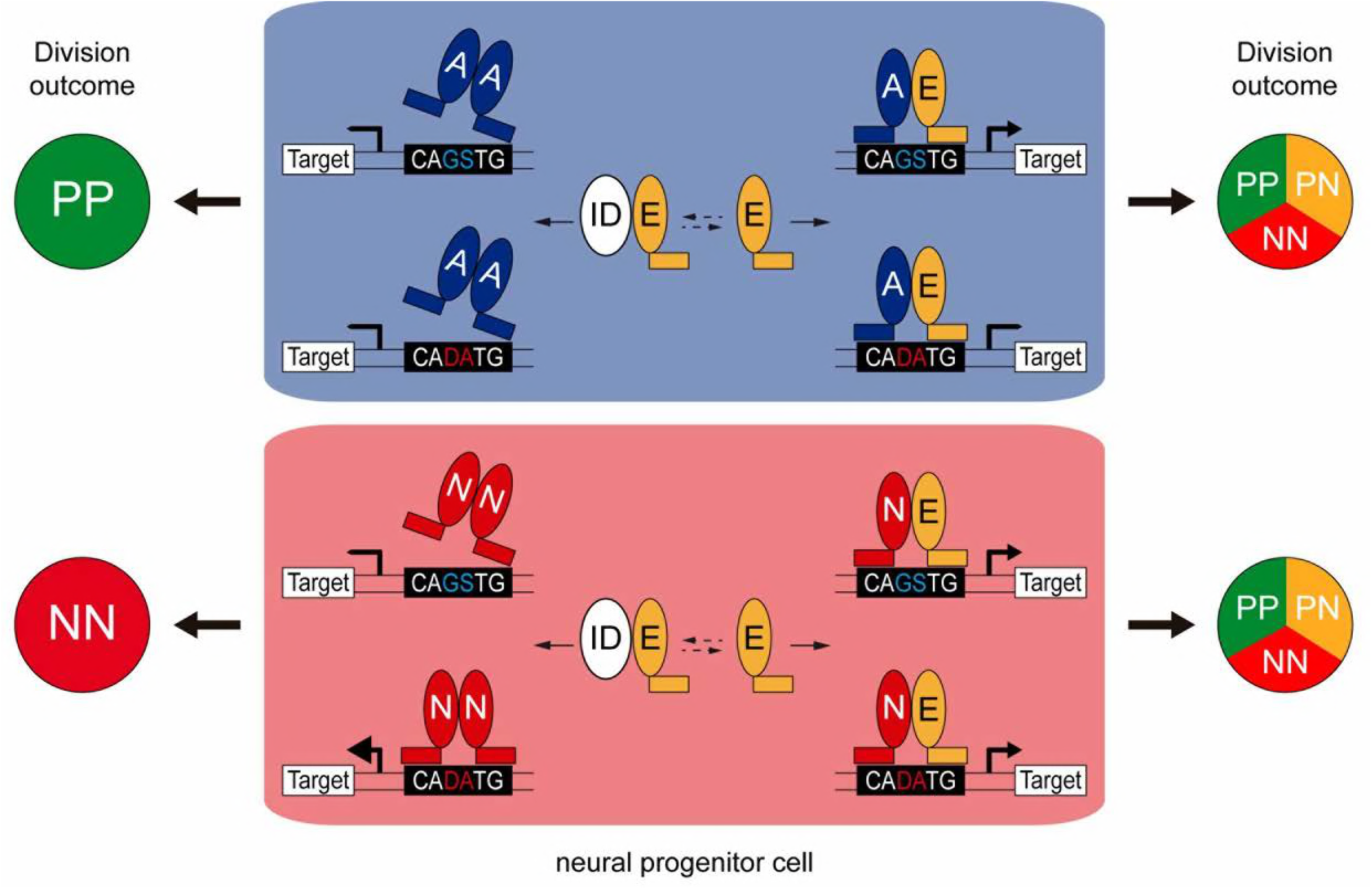
Model of the dual co-operation of E proteins with proneural proteins. In neural progenitors, ID proteins (ID) physically sequester E proteins (E), thereby regulating their ability to interact with ASCL1 and ATOH1 (A) or NEUROG1/2 (N). When E protein availability is limited, ASCL1/ATOH1 cannot bind optimally to high affinity CAGSTG E-box motifs, resulting in poor regulation of their target genes and favouring symmetric proliferative (PP) divisions and hence, progenitor maintenance. The release of E proteins from IDs allows heterodimerization with ASCL1/ATOH1, resulting in optimal binding to CAGSTG motifs, correct regulation of the target genes and the appropriate increase in neurogenic asymmetric (PN) and self-consuming (NN) divisions. In the absence of E proteins, NEUROG1/2 bind to high affinity CADATG motifs, possibly as homodimers, and regulate the expression of target genes in an exacerbated manner. This deregulation results in excessive neurogenic divisions that cause premature neuronal differentiation and depletion of the progenitor pool. In the presence of E proteins and when N-E heterodimers are formed, the activity of NEUROG1/2 is moderated and the proportions of the different modes of divisions are balanced appropriately to sustain the progenitor population while promoting correct neuronal differentiation.

On the one hand, our results support a revised model whereby E proteins synergize with proneural proteins specifically at CAGSTG E-boxes, the preferential motifs of E proteins (Lin et al, 2010; Pfurr et al, 2017). Therefore, E proteins facilitate the activity of the proneural proteins that share their preferential binding to CAGSTG motifs, such as ASCL1 and ATOH1 (Borromeo et al, 2014; Castro et al, 2011; Klisch et al, 2011; Lai et al, 2011). Inhibiting the activity of endogenous E proteins by expressing the E47bm mutant strongly impaired the generation of interneurons derived from spinal progenitors that express ATOH1 or ASCL1 alone. Conversely, enhancing the expression of E47 reinforced the ability of ATOH1 and more markedly, that of ASCL1 to promote neuronal differentiation. Our results suggest that this results from the capacity of E47 to increase the ability of these proneural proteins to trigger neurogenic divisions at the expense of proliferative ones (Figure 7). Such co-operation appears to be particularly crucial in the case of ASCL1, whose overexpression could barely increase neurogenic divisions *per se.* These observations support a growing body of evidence that ASCL1 possesses a mild neurogenic potential. For instance, the broad dP3-dP5 domain of spinal progenitors, in which ASCL1 is expressed alone or in combination with PTF1a, expands at the end of the first neurogenic wave before producing large numbers of dILA/B neurons (Borromeo et al, 2014; Wildner et al, 2006). Later on, ASCL1 is also involved in promoting oligodendrogenesis in both the developing brain and spinal cord (Huang et al, 2014). Moreover, recent studies have reported cell cycle promoting-genes among the targets bound by ASCL1 in the ventral telencephalon and that it also sustains the proliferation of adult neural stem cells (Castro et al, 2011; Urban et al, 2016), suggesting that its mild neurogenic ability might actually be required to sustain long-term production of the neural lineages. Whether the ability of ASCL1 to maintain neural progenitor pools is related to its dependence on the availability of E proteins is an intriguing hypothesis that would be worth testing.

On the other hand, our findings demonstrate that E proteins inhibit proneural protein binding to CADATG motifs. In consequence, E proteins restrict the activity of the proneural proteins that preferentially bind to these motifs, such as NEUROG1/2 (Madelaine & Blader, 2011; Seo et al, 2007). Since E47 restrains the capacity of NEUROG1/2 to promote neuronal differentiation in the context of both spinal neurogenesis and corticogenesis, this would appear to be a general feature of their behaviour. Early E47 depletion was recently shown to increase the production of both TBR1^+^ and SATB2^+^ neurons at mid-corticogenesis (Pfurr et al, 2017). In fact, the loss of E47 in early cortical progenitors, for which NEUROG2 constitutes the main proneural protein, causes premature neuronal differentiation. This is consistent with our model and it contrasts with the block in neuronal differentiation that would be expected if E47 was essential for NEUROG2 activity.

Our results also suggest that NEUROGs do not necessarily need to form heterodimers with E proteins to trigger neuronal differentiation. Indeed, NEUROG1/2-dependent differentiation is only mildly affected by the loss of E47 activity, and forced NEUROG1 homodimers more efficiently drive CADATG dependent transcription and neuronal differentiation than NEUROG1-E47 heterodimers. Similarly, NEUROG2 homodimers better transactivate neuronal differentiation genes than NEUROG2-E47 heterodimers (Li et al, 2012), and EMSA experiments suggest that multiple combinations of proneural homo- and heterodimers exist (Henke et al, 2009). The physiological relevance of such proneural homodimers is worthy of further study but to date, our attempts to determine whether NEUROG1 homodimers are formed *in vivo* during spinal neurogenesis remain inconclusive for technical reasons (data not shown). Nevertheless, the strong capacity of NEUROGs to trigger neurogenic divisions independent of E proteins, including self-consuming NN divisions, correlates well with the fact that neural progenitors expressing NEUROG1/NEUROG2 are usually depleted during the neurogenic phase (Kim et al, 2011; Simmons et al, 2001). Together, these results support the notion that E proteins are required to dampen the capacity of NEUROGs to trigger neurogenic divisions, thereby avoiding the premature depletion of neural progenitor pools (Figure 7).

This dual mode of action of E proteins in conjunction with ASCL1/ATOH1 or NEUROG1/NEUROG2 would also explain why modulating canonical BMP activity affects differently the generation of the distinct neuronal subtypes produced during primary spinal neurogenesis. Inhibiting BMP7 or SMAD1/5 would result in the release of E proteins from their complexes with IDs. In turn, this would facilitate ATOH1 and ASCL1 activity, prematurely increasing the proportion of neurogenic divisions undertaken by the corresponding dP1 and dP3/dP5/p2 progenitors, causing their premature differentiation and exhaustion, and ultimately leading to a production of fewer neurons. As NEUROGs are less dependent on E proteins, the inhibition of canonical BMP signalling only mildly impairs the generation of the neuronal subtypes that derive from progenitors expressing NEUROG1/NEUROG2.

In summary, the results presented here led us to propose that E proteins fine-tune neurogenesis by buffering the activity of the distinct proneural proteins. As such, these data add another layer of sophistication to the molecular mechanisms that regulate the activity of proneural bHLH proteins and hence, neurogenesis.

## Materials and Methods

### In ovo electroporation

Fertilized white Leghorn chicken eggs were provided by Granja Gibert, rambla Regueral, S/N, 43850 Cambrils, Spain. Eggs were incubated in a humidified atmosphere at 38·C in a Javier Masalles 240N incubator for the appropriate duration and staged according to the method of Hamburger and Hamilton (HH, (Hamburger & Hamilton, 1951). According to EU animal care guidelines, no IACUC approval was necessary to perform the experiments described herein, considering that the embryos used in this study were always harvested at early stages of embryonic development (at E5 at the latest). Sex was not identified at these stages.

Unilateral *in ovo* electroporations in the developing chick spinal cord and dorsal telencephalon were performed respectively at stages HH14-15 and HH18 (54 and 69 hours of incubation). In the telencephalon, corticogenesis was studied specifically in the dorsal-medial-lateral (dML) subregion to minimize any possible variability along the medial-lateral axis. Plasmids were diluted in RNAse-free water at the required concentration [0 to 4 μg/μl] and injected into the lumen of the caudal neural tube or the right cerebral ventricle using a fine glass needle. Electroporation was triggered by applying 5 pulses of 50 ms at 22.5 V with 50 ms intervals using an Intracel Dual Pulse (TSS10) electroporator. Electroporated chicken embryos were incubated back at 37C and recovered at the times indicated (16-48 hours post-electroporation).

### Plasmids

To facilitate comparisons in gain-of-function experiments, all the constructs used in this study were inserted under the control of a pCAGGS promoter that harbors high activity in chick (pCAGGS or pCAGGS_ires_GFP, kindly provided by Andy McMahon, Megason & McMahon, 2002), and were electroporated at similar concentrations (0, 0.1, 0.5 or 1 μg/μl as specified in the respective figure legends). Non-fluorescent pCAGGS plasmids were combined with 0.25 μg/μl of pCS2_H2B-GFP for visualization. The pCAGGS:ASCL1, pCAGGS:NGN1 and pCAGGS:NGN2 plasmids were kindly provided by François Guillemot. The pCAGGS:ATOH1_ires_GFP plasmid was obtained by subcloning from a pCMV:ATOH1 kindly provided by Nissim Ben-Arie (Krizhanovsky et al, 2006). The pCAGGS:E47 and pCAGGS:TCF12 were kindly provided by Jonas Muhr (Holmberg et al, 2008). The pCAGGS:E47bm_ires_GFP plasmid was derived from a pGK:E47_CFP plasmid kindly provided by Yuan Zhang (Zhuang et al, 1998). The pCAGGS:ID2_ires_GFP was derived from a pCMV:ID2 plasmid, and the pCAGGS_SMAD5-SD_ires_GFP was described previously (Le Dreau et al, 2012). Only Somitabun (pCS2:Somitabun, kindly provided by Jonathan Slack, Beck et al, 2001) and NGN1-AQ and its wild-type NGN1 version (pMiW:myc-NGN1 and pMiW:myc-NGN1-AQ, kindly gifted by Jane Johnson, Gowan et al, 2001) were used in a different backbone. HA-tagged versions of ASCL1 (pCAGGS:HA-ASCL1, Alvarez-Rodriguez & Pons, 2009) and NGN1 (pCAGGS:HA-NGN1) and a pCMV2_Flag-E47-RFP plasmid kindly provided by Yoshihiro Yoneda (Mehmood et al, 2009) were used for chromatin immuneprecipitation assays. Inhibition of cBmp7, cSmad1, cSmad5 or cId2 expression was triggered by electroporation of short-hairpin constructs inserted into the pSuper (Oligoengine) or pSHIN (Kojima et al, 2004) vectors. Electroporation of 2-4 μg/μl of these constructs caused a specific and reproducible 50% inhibition of the target expression (see Le Dreau et al, 2012). The pSox2:GFP and pTis21:RFP reporters used to assess the modes of divisions undergone by spinal progenitors were previously described in details (Saade et al, 2013). The pSox2:luc derived from the pSox2:GFP (Saade et al, 2013). The different versions of the pId2:luc reporters were kindly provided by Yoshifumi Yokota (Kurooka et al, 2012). The pkE7:luc (Akazawa et al, 1995) and pNeuroD:luc reporters were kindly provided by Masashi Kawaichi and François Guillemot, respectively. The pNeuroDmut:luc reporter was obtained by site-directed mutagenesis of the single CAGGTG E-box contained in the NeuroD promoter region. The pTubb3:luc reporter was obtained by subcloning the Tubb3 enhancer region present in the pTubb3enh:GFP plasmid kindly provided by Jonas Muhr (Bergsland et al, 2011) into the pGL3:luc vector (Promega).

### Generation of tethered constructs

The tethered bHLH dimers were derived from the pCAGGS:ASCL1-t-E47_ires_GFP kindly provided by François Guillemot (Geoffroy et al, 2009). This plasmid and pCAGGS:NGN1 were used as templates to generate the pCAGGS:ASCL1-t-ASCL1_ires_GFP, pCAGGS:NGN1-t-E47_ires_GFP and pCAGGS:NGN1-t-NGN1_ires_GFP plasmids, using a tether peptide AAAGTSAGGAAAGTSASAATGA flanked by SpeI and ClaI restriction sites as described previously (Henke et al, 2009). Expression of the tethered bHLH dimers was assessed by western blot after transfection into HEK293 cells. Transient cell transfections were obtained by electroporation applying 2 pulses of 120V, 30ms (Microporator MP-100, Digital Bio). Cells were grown for 24 hours onto poly-L-Lysine-coated 6-well dishes in DMEM/F12 supplemented with 10% fetal bovine serum and 50 mg/L of Gentamicin until reaching 70-80% confluence. The typical transfection efficiency of this procedure was 40-60%. Cells were lysed in 1X SDS loading buffer (10% glycerol, 2% SDS, 100 mM dithiothreitol, and 62.5 mM Tris-HCl, pH 6.8) and DNA was disrupted by sonication. Protein extracts were separated by SDS-PAGE electrophoresis, transferred to Immobilon-FL PVDF membranes (IPFL00010, Millipore), blocked with the Odyssey Blocking Buffer (927-40000, LI-COR), and incubated with antibodies against ASCL1 (BD Pharmingen, cat#556604, 1:1000), NGN1 (Millipore, cat#AB15616, 1:3000) or E2A (Santa Cruz, cat#sc-763, 1:1000). Detection was performed using fluorescence-conjugated secondary antibodies and an Odyssey Imaging System (LICOR).

### Immunohistochemistry

For immunohistochemistry experiments, chicken embryos were carefully dissected, fixed for 2 hours at 4°C in 4% paraformaldehyde and rinsed in PBS. Immunostaining was performed on either vibratome (40 μm) or cryostat (16 μm) sections following standard procedures. After washing in PBS-0.1% Triton, the sections were incubated overnight at 4C with the appropriate primary antibodies (Supplementary File l) diluted in a solution of PBS-0.1% Triton supplemented with 10% bovine serum albumin or sheep serum. After washing in PBS-0.1% Triton, sections were incubated for 2 hours at room temperature with the appropriate secondary antibodies diluted in a solution of PBS-0.1% Triton supplemented with 10% bovine serum albumin or sheep serum. Alexa488-, Alexa555- and Cy5-conjugated secondary antibodies were obtained from Invitrogen and Jackson Laboratories. Sections were finally stained with 1 μg/ml DAPI and mounted in Mowiol (Sigma-Aldrich).

### Image acquisition, treatment and quantification

Optical sections of fixed samples (transverse views of the spinal cord, coronal views for the telencephalon) were acquired at room temperature with the Leica LAS software, in a Leica SP5 confocal microscope using 10x (dry HC PL APO, NA 0.40), 20x (dry HC PL APO, NA 0.70), 40x (oil HCX PL APO, NA 1.25-0.75) or 63x (oil HCX PL APO, NA 1.40-0.60) objective lenses. Maximal projections obtained from 2μm Z-stack images were processed in Photoshop CS5 (Adobe) or ImageJ for image merging, resizing and cell counting.

Quantification of endogenous ID2 intensity was assessed using the ImageJ software. Cell nuclei of H2B-GFP^+^ electroporated and neighboring non-electroporated cells were delimitated by polygonal selection, and the mean intensity of ID2 immunoreactivity quantified as mean gray values. Quantifications were performed on at least six electroporated and six non-electroporated cells per image, in at least three different images per embryo.

### In situ hybridization

Chicken embryos were recovered at the indicated stage, fixed overnight at 4°C in 4% PFA, rinsed in PBS and processed for whole mount RNA in situ hybridization following standard procedures. Probes against chick cId2 (#chEST852M19) and cNgn2 (#chEST387d10) were purchased from the chicken EST project (UK-HGMP RC). Probes against cTcf3/E2a, cAscl1 and cNgn1 were kindly provided by Drs Jonas Muhr, José-Maria Frade and Cristina Pujades. The probe against cTcf12/cHeb was obtained by PCR from genomic DNA of E4 chicken embryonic tissue and the purified 623 nucleotides insert was sub-cloned into the pGEM-T vector (Promega). Hybridized embryos were post-fixed in 4% PFA and washed in PBT. 45μM-thick sections were cut with a vibratome (VT1000S, Leica), mounted and photographed using a microscope (DC300, Leica). The data show representative images obtained from 3 embryos for each probe.

### Luciferase assay

Transcriptional activity was assessed following electroporation of the reporters pkE7:luc (gift from Masashi Kawaichi), pNeuroD:luc (gift from François Guillemot), pNeuroDmut:luc, pSox2:luc or the different versions of pId2:luc (provided by Yoshifumi Yokota) together with a renilla luciferase reporter used for normalization and the indicated bHLH TF-encoding plasmids. Embryos were harvested 24 hours later and GFP-positive neural tubes were dissected and homogenized in a Passive Lysis Buffer on ice. Firefly- and renilla-luciferase activities were measured by the Dual Luciferase Reporter Assay System (Promega).

### Cell cycle exit assay

The average number of divisions undergone by electroporated spinal progenitors was assessed *in vivo* using the CellTrace Violet Cell Proliferation Kit (Invitrogen). The Violet cell tracer (1 mM), a cytoplasmic retention dye that becomes diluted as cells divide, was injected into the lumen of the neural tube at the time of electroporation. Embryos were recovered 48 hours later, the neural tubes were carefully dissected and recovered and the cells dissociated following a 10-15 min digestion in Trypsin-EDTA (Sigma). The fluorescence intensity of the Violet tracer was measured in viable dissociated electroporated GFP^+^ cells in the 405/450nm excitation/emission range on a Gallios flow cytometer (Beckman Coulter, Inc).

### Assessment of the modes of divisions

Chicken embryos were recovered 16 hours after co-electroporation of the pSox2:eGFP and pTis21:RFP reporters together with the indicated bHLH TF-encoding plasmids. Cell suspensions were obtained from pools of 6-8 dissected neural tubes after digestion with Trypsin-EDTA (Sigma) for 10-15 min, and further processed on a FACS Aria III cell sorter (BD Biosciences) for measurement of eGFP and RFP fluorescences. At least 1,000 cells for each progenitor population (PP, PN and NN) were analyzed per sample.

### Chromatin immunoprecipitation assay

HEK293 cells were transfected by a standard calcium phosphate co-precipitation protocol with combinations of pCAGGS_ires_GFP, pCAGGS:HA-ASCL1, pCAGGS:HA-NGN1, pCMV2_Flag-E47-RFP together with the pkE7:luc or pNeuroD:luc reporters, with a total of 10 μg of DNA per 100 mm dish. 24 hours later, cells were collected and 10% of the material was reserved to check transfection by Western blot. For chromatin immunoprecipitation assays, approximately 1 million trasnfected HEK293 cells were fixed with 1% formaldehyde for 10 minutes at room temperature. Fixation was quenched by adding 0.125M glycine for 5 minutes. After 2 washes with PBS, cells were lysed on ice for 20 minutes in a lysis buffer containing protease inhibitors (1% SDS; 10mM EDTA pH8.0; 50mM Tris-HCl pH8.1). Sonication was performed with a Bioruptor sonicator to obtain 200-500bp shredded chromatin fragments. Chromatin purification was carried out by spinning samples down at maximum speed at 4C during 30 minutes. Purified chromatin was pre-cleared with protein A agarose (Millipore #16-125) for 30 minutes. 25 μg of chromatin were immunoprecipitated with 5μL of anti-RFP serum (Herrera et al, 2014), 2μg of anti-HA (Abcam, cat#20084), anti-NGN1 (Millipore, cat#15616) or unspecific rabbit IgG (Diagenode, cat#C15410206) antibodies. Antibody-chromatin complexes were recovered using magnetic beads (Magna ChIP, Millipore, cat#16-661) and immuno-complexes were washed once with TSE I (0.1% SDS; 1% Triton-X100; 2mM EDTA pH8.0; 20mM Tris-HCl pH8.1; 150mM NaCl), TSE II (0.1% SDS; 1% Triton-X100; 2mM EDTA pH8.0; 20mM Tris-HCl pH8.1; 500mM NaCl), TSE III (0.25M LiCl; 1% NP-40; 1% Sodium Deoxicholate; 1mM EDTA pH8.0; 10mM Tris-HCl pH8.1) and twice with TE (Tris-HCl 10mM, EDTA 1mM). Reversal of crosslinking was done by incubating samples in elution buffer (1% SDS, 0.1M NaHCO3) overnight at 65C. DNA was purified by phenol-chloroform extraction followed by ethanol precipitation. Quantification of the DNA target regions and negative control (luciferase ORF) was assessed by qPCR in a Lightcycler 480 (Roche) using specific primers (Supplementary File 2).

Proteins extracts were obtained by incubation in a RIPA buffer (150 mM NaCl, 1.0% NP-40,0.5% sodium deoxycholate, 0.1% SDS and 50 mM Tris, pH 8.0) supplemented with protease and phosphatase inhibitors for 20 minutes on ice and centrifugation (20 minutes at maximum speed). 30 μg of protein samples were mixed with the Laemmli buffer (375 mM Tris pH =6.8, 12%SDS, 60% glycerol, 600 mM DTT, 0.06% bromphenol blue), heated to 95C and then separated on a SDS-PAGE gel in running buffer (25 mM Tris base, 190 mM glycine, 0.1% SDS, pH=8,3). Proteins were transferred to a nitrocellulose membrane using transfer buffer (190 mM glycine, 25mM Tris, 20% Methanol, 0.1% SDS) for 90 minutes at 80V. Membranes were blocked for 1 h with a solution of PBS-5% milk, 1% Tween (PBST) and further incubated overnight at 4C with appropriate primary antibodies diluted in PBST: rabbit anti-HA (Abcam, cat #ab20084), rabbit anti-RFP serum (Herrera et al, 2014) and mouse anti-Tubulin beta (Millipore, cat #MAB3408). After three washes in PBST, membranes were incubated with Horseradish peroxidase-conjugated anti-rabbit IgG or anti-mouse IgG secondary antibodies (Sigma-Aldrich, cat#GENA934-1ML and cat#GENA931) for 1 hour at room temperature and the signals detected by chemiluminescence using the Immobilon western chemiluminiscent HRP substrate (Sigma-Aldrich, cat# WBKLS0100).

### Statistical analyses

No statistical method was used to predetermine sample size. The experiments were not randomized. The investigators were not blinded to allocation during experiments or outcome assessment. Statistical analyses were performed using the GraphPad Prism 6 software (GraphPad Software, Inc.). For *in vivo* experiments, cell counts were typically performed on 2-5 images per embryo and *n* values correspond to different embryos, except for the assessment of the modes of divisions where *n* values correspond to pools of embryos. For *in vitro* chromatin immunoprecipitation assays, *n* values represent the numbers of independent experiments performed. The *n* values are indicated in the corresponding figure legend. The normal distribution of the values was assessed by the Shapiro-Wilk normality test. Significance was then assessed with a two-sided unpaired t-test, one-way ANOVA + Tukey’s test or two-way ANOVA + Sidak’s test for data presenting a normal distribution, or alternatively with non-parametric Mann–Whitney or Kruskal-Wallis + Dunn’s multiple comparisons’ tests for non-normally distributed data. n.s: non-significant; ^*^: p<0.05 or less, as indicated in individual figures.

## Acknowledgments

We thank the members of the laboratory for helpful comments on the manuscript. We are grateful to Drs N. Ben-Arie, C. Birchmeier, J-M. Frade, F. Giraldez, F. Guillemot, T.M. Jessell, J.E. Johnson, A. Joyner, M. Kawaichi, A McMahon, J. Muhr, T. Müller, S. Pfaff, C. Pujades, J. Slack, Y. Yokota, Y. Yoneda and Y. Zhuang for kindly providing reagents. We also thank the Developmental Studies Hybridoma Bank, developed under the auspices of the NICHD and maintained by The University of Iowa, Department of Biological Sciences, Iowa City, IA, USA. We acknowledge E. Rebollo and the IBMB Molecular Imaging platform and J. Comas and the PCB Flow Cytometry facility for excellent assistance. This work was supported by the grants to E.M from BFU2016-81887-REDT and BFU2016-77498-P. G. LD was supported by AECC (AIO2014). R.E and A. H were recipient of postdoctoral fellowships from the Mexican National Council of Science and Technology (CONACYT) and Spanish Ministry of Education, Industry and Competitiveness (MINECO, Program Juan de la Cierva, #FJCI-2015-26175). R.F was recipient of a Ph.D fellowship from the Spanish Ministry of Education, Culture and Sports (MECD, program FPU, #FPU13/01384).

## Additional Files

- Supplementary File 1: List of antibodies
- Supplementary File 2: List of primers used for ChIP assays.

**Figure 1-figure supplement 1:**
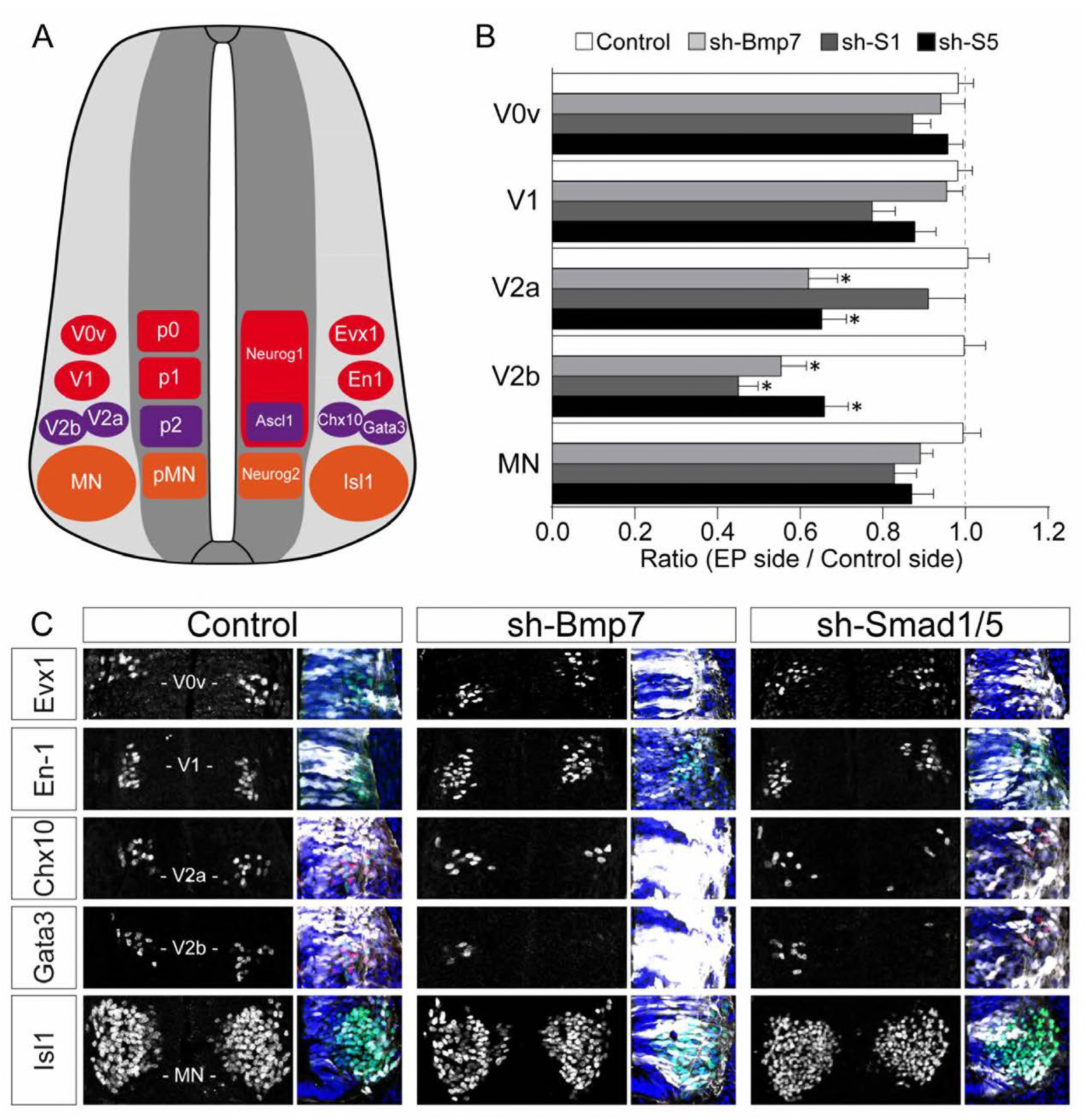
Inhibiting the canonical BMP pathway affects the generation of ventral spinal neurons. (A) Diagram of a transverse section of the developing spinal cord at mid-neurogenesis, highlighting the ventral neuron subtypes analysed and the markers used to identify them. (B, C) The proportions (B) and representative images (C) of the ventral spinal neuron subtypes generated 48 hours after in ovo electroporation with a control plasmid or plasmids producing sh-RNAs specifically targeting cBmp7 (sh-Bmp7), cSmadl (sh-Smad1) or cSmadS (sh-Smad5). GFP staining (white) and DARI (blue) are shown to confirm the region of interest was electroporated. The data are presented as the mean ratios ± sem obtained from n=6-17 embryos per condition; one-way ANOVA + Tukey’s test; ^*^P<0 05

**Figure 2-figure supplement 1:**
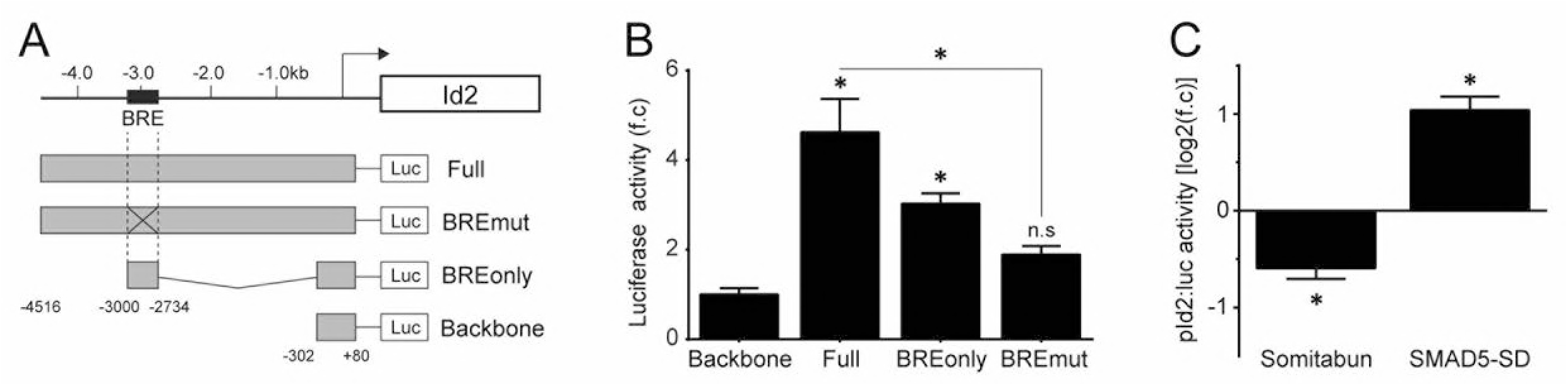
Regulation of ID2 expression by the canonical BMP pathway. (A) Representation of the proximal murine *Id2* promoter region and different mutant constructs of pld2:luc reporters, highlighting the location of the SMAD1/5/8-responsive (BRE) elements. (B) Transcriptional assay showing the activity of the different pld2:luc reporters measured 24 hpe. The data are expressed as the mean fold change ± sem relative to the control values, obtained from n=7-8 embryos per condition; one-way ANOVA + Tukey’s test; ^*^P<0.05. (C) Transcriptional assay showing the activity of the full pld2:luc reporter measured 24 hpe with dominant-negative (Somitabun) or constitutively active (SMAD5-SD) SMAD5 mutant constructs. The data are expressed as the mean Log2 fold changes ± sem relative to the control values, obtained from n=7-10 embryos; two-sided unpaired t-test; ^*^P<0.05.

**Figure 2-figure supplement 2:**
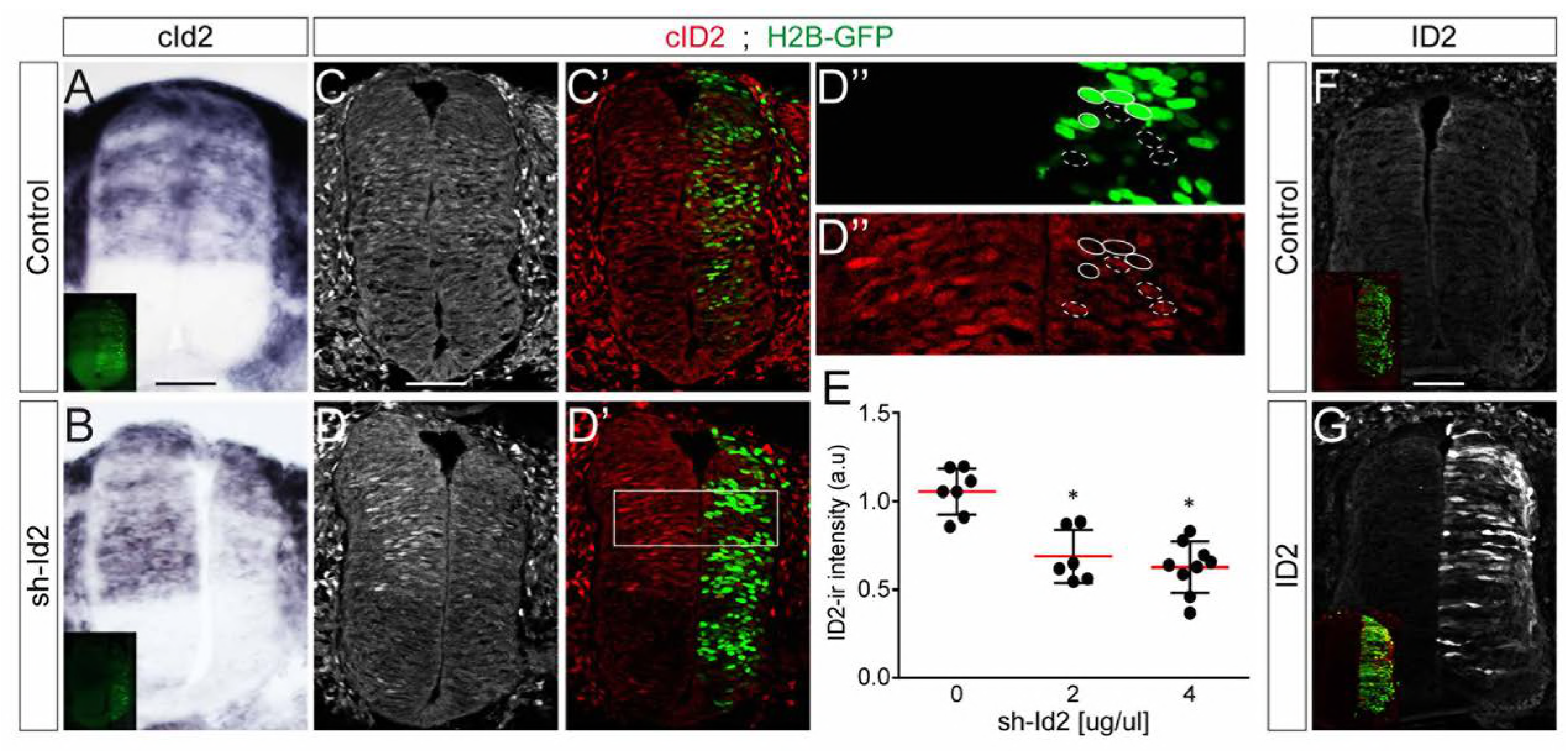
Modulation of ID2 activity *in vivo.* (A-B) Detection of cld2 transcripts by in situ hybridization in transverse spinal sections 24 hpe with control (A) or sh-ld2-producing (B) plasmids. (C-D) Endogenous clD2 immunoreactivity detected in transverse spinal sections 24 hpe with control (C, C’) or sh-!d2-producing (D-D”) plasmids, and quantified in electroporated and nearby non-electroporated cells (as highlighted in D”). (E) The data represent the mean clD2 immunoreactivity ± sd measured after electroporation of a control plasmid [0] or increasing concentrations [2 and 4 μg/μl] of sh-ld2 plasmids in electroporated relative to non-electroporated cells, obtained from n=6-9 embryos per condition; one-way ANOVA + Tukey’s test; ^*^P<0.05. (F-G) ID2 immunoreactivity in transverse spinal sections 24 hpe with a control plasmid (F) or overexpression of a murine ID2 construct (G). Scale bars, 50 μM.

**Figure 3-figure supplement 1:**
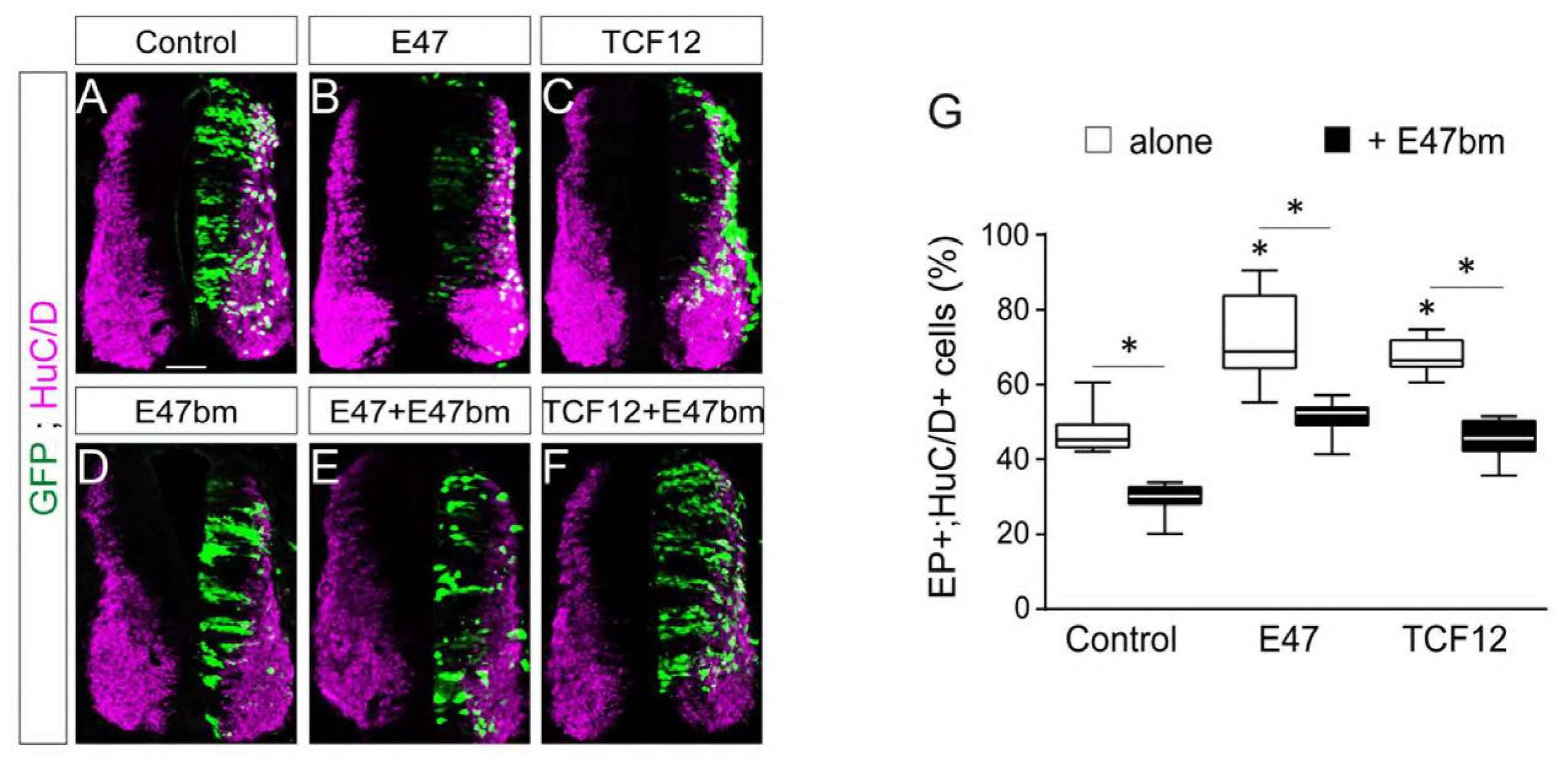
E47bm rescues the premature neuronal differentiation caused by both E47 and TCF12. (A-F) Transverse spinal cord sections of electroporated cells (GFP+) that had differentiated into neurons (HuC/D+) 48 hpe with a control (A), E47 (B), TCF12 (C), E47bm (D) construct or combinations thereof (E, F). (G) The box-and-whisker plots show the proportion obtained from n=7-14 embryos per condition; one-way A NOVA + Tukey’s test; ^*^P<0.05. Scale bars, 50 μM

**Figure 4-figure supplement 1:**
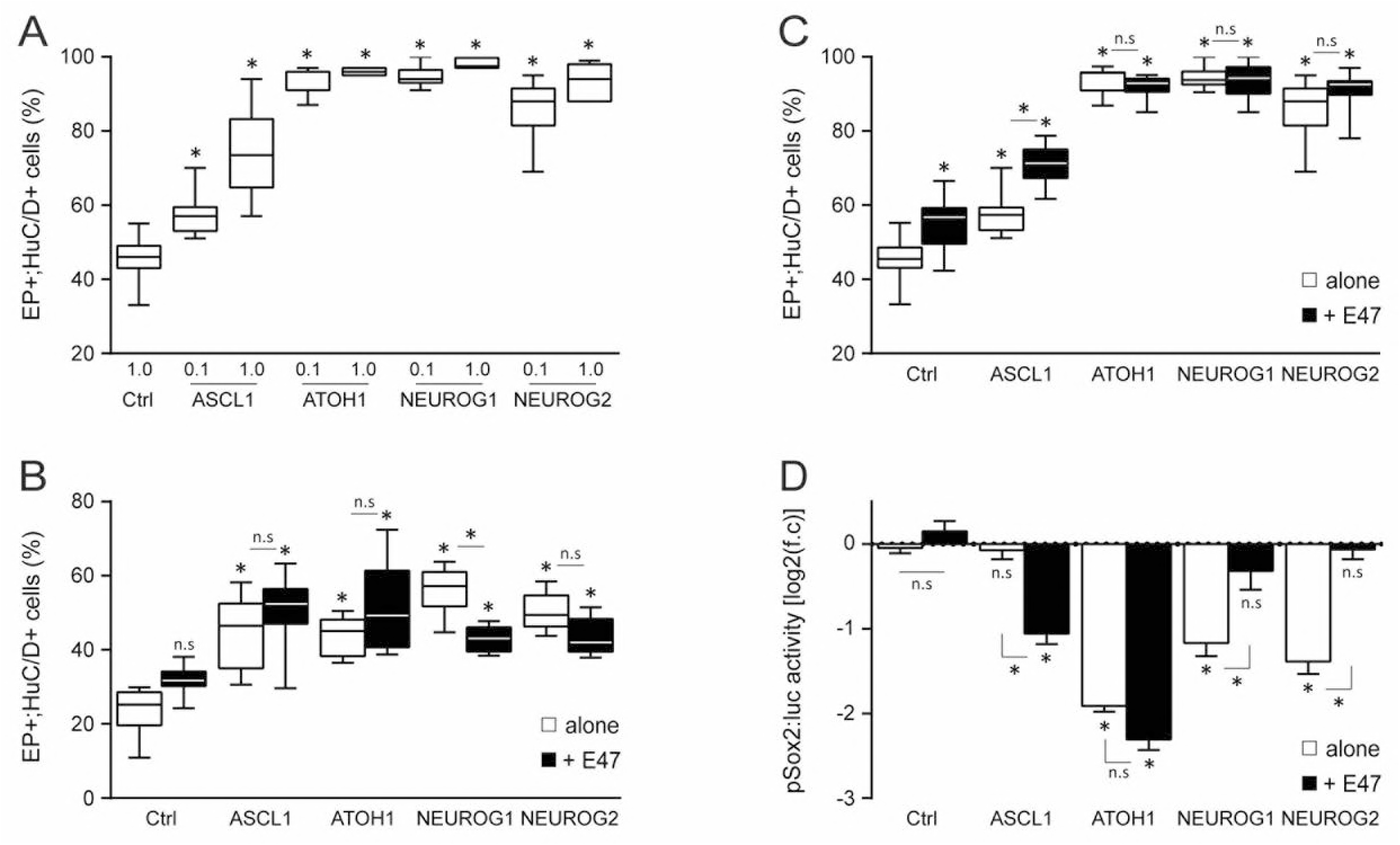
Effects of E47 and proneural proteins on spinal neuronal differentiation. (A) Box-and-whisker plots showing the proportion of electroporated cells (GFP+) that differentiated into neurons (HuC/D+) 48 hpe with increasing concentrations [0.1 or 1 μg/μl] of ASCL1, ATOH1, NEUROG1 or NEUROG2, from n=7-13 embryos per condition; one-way ANOVA + Tukey’s test; ^*^P<0.05. (B, C) Proportion of electroporated cells (GFP+) that differentiated into neurons (HuC/D+) 24 (B) or 48 (C) hpe with control, ASCL1, ATOH1, NEUROG1 or NEUROG2 alone (white whiskers) or together with E47 (black whiskers), obtained from n=6-13 embryos; one-way ANOVA + Tukey’s test; ^*^P<0.05. (D) Transcriptional assay showing the activity of a pSox2:luc reporter measured 24 hpe with control, ASCL1, ATOH1, NEUROG1 or NEUROG2 alone (white whiskers) or together with E47 (black whiskers). The data are expressed In Log2 as the mean fold change + sem relative to the control values, obtained from n=6-19 embryos per condition; one-way ANOVA + Tukey’s test; ^*^P<0.05.

**Figure 5-figure supplement 1:**
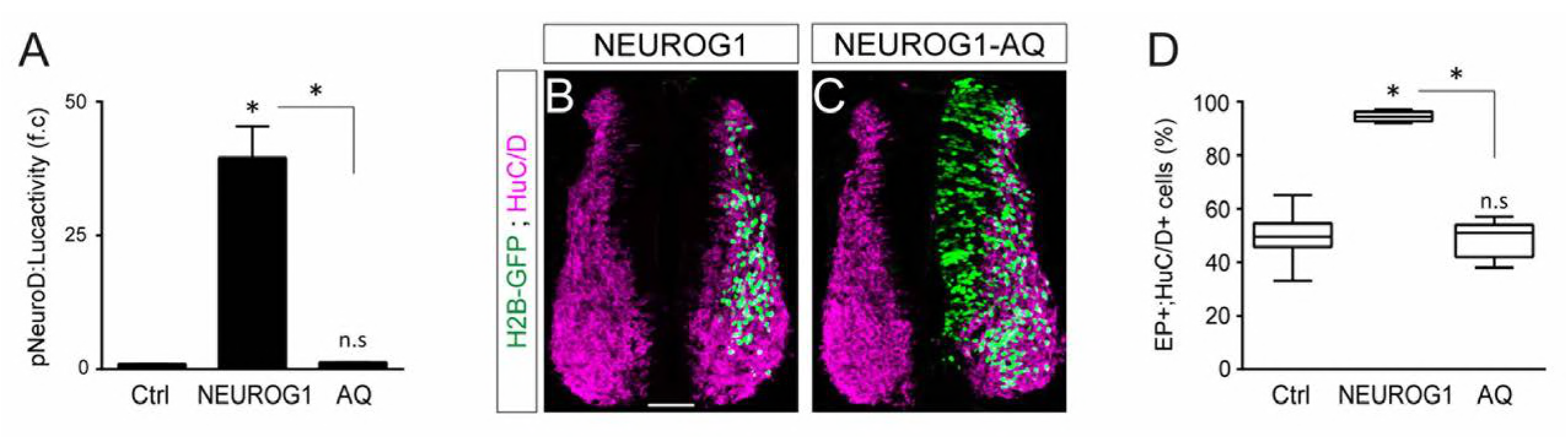
The ability of NEUROG1 to induce spinal neuronal differentiation depends on its ONA-binding. (A) Transcriptional assay showing the activity of the pNeuroD luc reporter measured 24 hpe with control or myc-tagged wild-type NEUROG1 construct and the NEUROG1-AQ mutant, obtained from n=6 embryos per condition; Kruskal-Wallis + Dunn’ test; ^*^P<0.05. (B, C) Transverse spinal cord sections of electroporated cells (H2B-GFP+) that differentiated into neurons (HuC/D+) 48 hpe with NEUROG1 (B) or NEUROG1-AQ (C). (D) Box-and-whisker plots showing the proportion obtained from n=6-7 embryos per condition; Kruskal-Wallis + Dunn’ test; ^*^P<0 05 Scale bars, 50 μM

**Figure 5-figure supplement 2:**
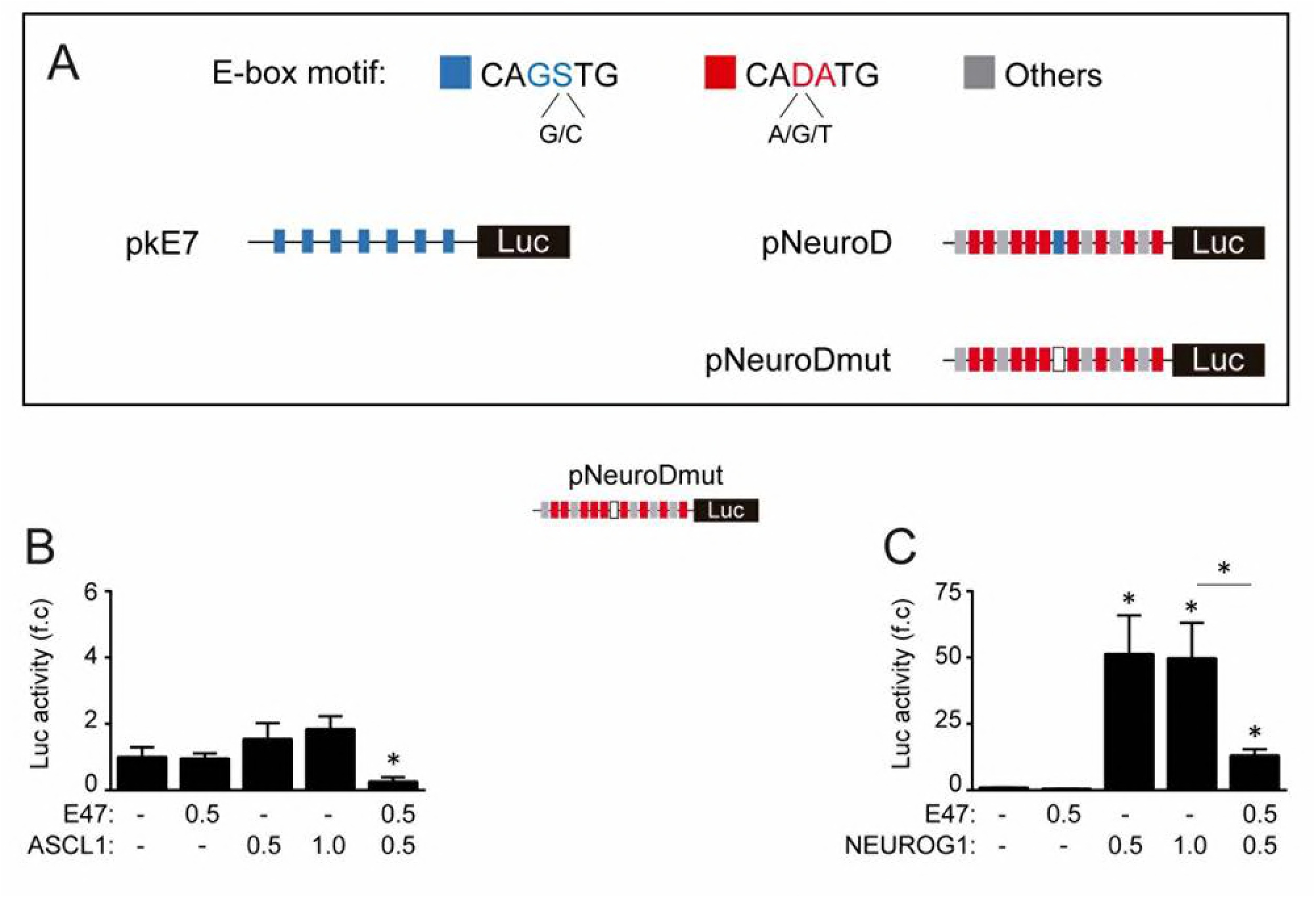
E-box dependent activity of E47, ASCL1 and NEUR0G1 during spinal neurogenesis. (A) Schematic representation of the E-box containing luciferase reporters used in this study. (B, C) Transcriptional assays showing the activity of the mutated version of the pNeuroD:luc reporter measured 24 hpe with control, E47 and ASCL1 (B) or NEUROG1 (C). The data are expressed as the mean fold change ± sem relative to the control values, obtained from n=7-13 (B) or 6 (C) embryos per condition; Kruskal-Wallis + Dunn’ test or one-way ANOVA + Tukey’s test; ^*^P<0.05.

**Figure 5-figure supplement 3:**
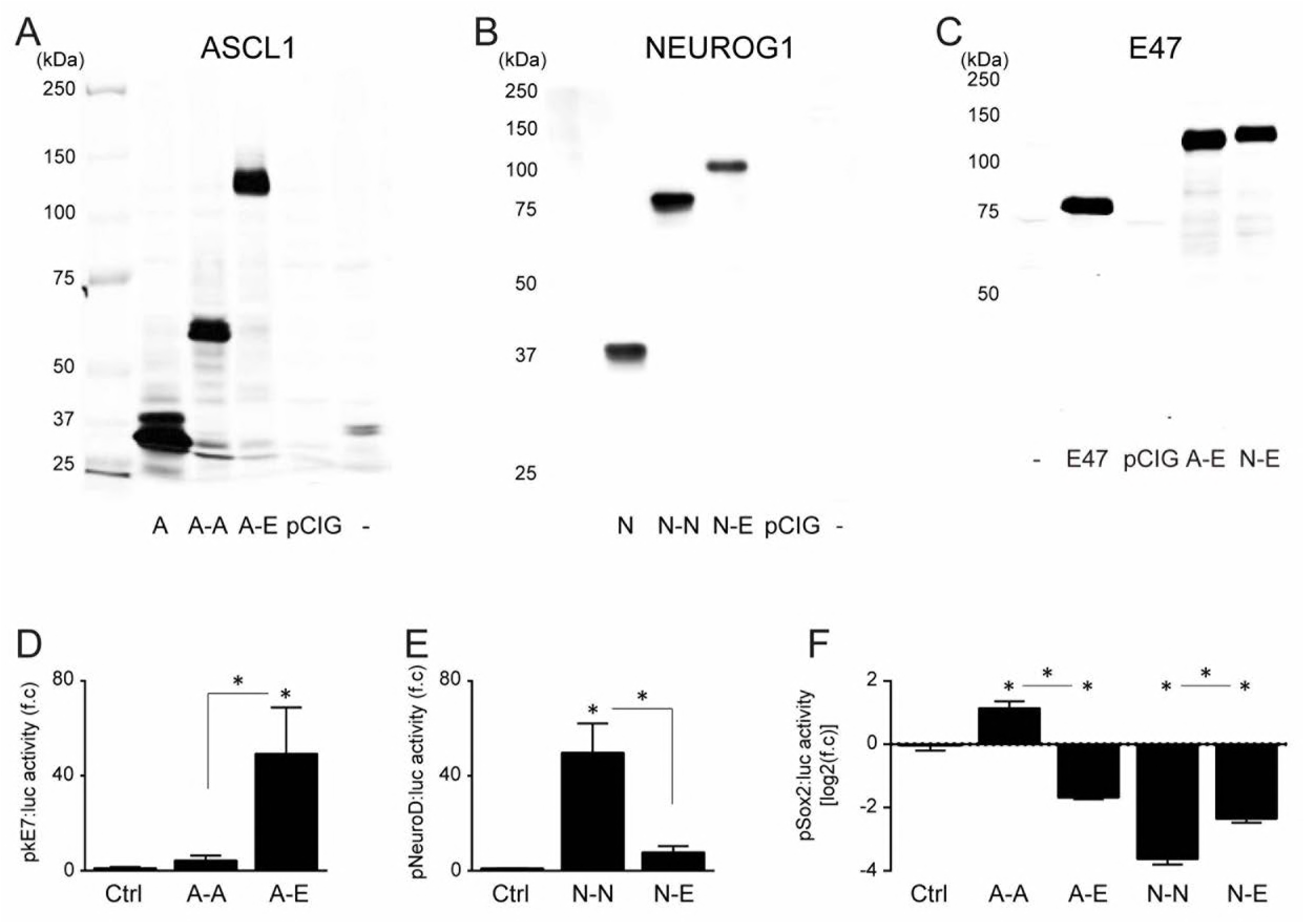
Characterization of the tethered constructs of bHLH dimers. (A-C) Western blot detection of monomeric and dimeric ASCL1 (A), NEUROG1 (B) and E47 (C) constructs in protein extracts obtained 24 hours after transfecting HEK293 cells with constructs encoding ASCL1, NEUROG1 and E47 monomers (A, N, E), homo- and heterodimers (A-A, A-E, N-N, N-E), a control plasmid (pCIG), or non-transfected cells (-). (D-F) Transcriptional assays show the activity of the pkE7 (D), pNeuroD (E) and pSox2 (F) luciferase reporters measured 24 hpe with controls, ASCL1 (A-A) and NEUROG1 (N-N) homodimers, and ASCL1-E47 (A-E) and NEUROG1-E47 (N-E) heterodimers. The data are expressed as the mean (D, E) or Log2 fold changes (F) ± sem relative to the control values, obtained from n=4-6 (D), 11-12 (E) or 6 (F) embryos per condition; Kruskal-Wallis + Dunn’ test or one-way ANOVA + Tukey’s test; ^*^P<0.05.

**Figure 6-figure supplement 1:**
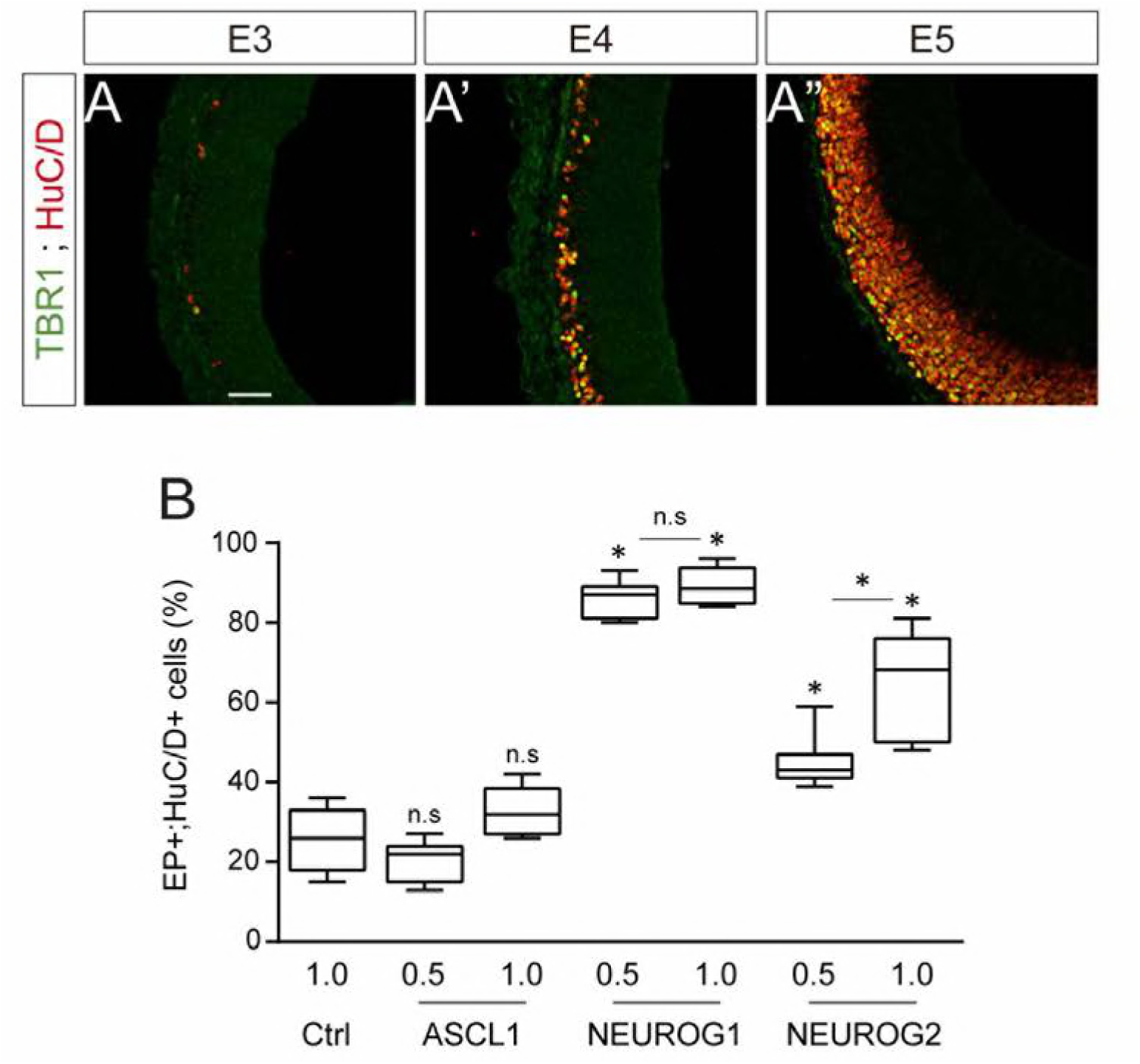
Neurogenesis and concentration dependent effects of proneural proteins during early chick corticogenesis. (A) Coronal telencephalic sections showing TBR1 immunoreactivity in differentiating neurons (HuC/D+) that are generated during early corticogenesis in chick, at E3 (A), E4 (A’) and E5 (A”) (B) Proportion of electroporated cells (GFP+) that differentiated Into neurons (HuC/D+) 48 hpe with increasing concentrations [0.5 or 1 μg/μl] of the ASCL1, NEUROG1 or NEUROG2 constructs. Data were obtained from n=6-9 embryos per condition; one-way ANOVA + Tukey’s test; ^*^P<0,05. Scale bar, 50 μM.

